# Accelerating Ligand Discovery for Insect Odorant Receptors

**DOI:** 10.1101/2024.09.12.612620

**Authors:** Arthur Comte, Maxence Lalis, Ludvine Brajon, Riccardo Moracci, Nicolas Montagné, Jérémie Topin, Emmanuelle Jacquin Joly, Sébastien Fiorucci

## Abstract

Odorant receptors (ORs) are main actors of the insects peripheral olfactory system, making them prime targets for pest control through olfactory disruption. Traditional methods employed in the context of chemical ecology for identifying OR ligands rely on analyzing compounds present in the insect’s environment or screening molecules with structures similar to known ligands. However, these approaches can be time-consuming and constrained by the limited chemical space they explore. Recent advances in OR structural understanding, coupled with scientific breakthroughs in protein structure prediction, have facilitated the application of structure-based virtual screening (SBVS) techniques for accelerated ligand discovery. Here, we report the first successful application of SBVS to insect ORs. We developed a unique workflow that combines molecular docking predictions, *in vivo* validation and behavioral assays to identify new behaviorally active volatiles for non-pheromonal receptors. This work serves as a proof of concept, laying the groundwork for future studies and highlighting the need for improved computational approaches. Finally, we propose a simple model for predicting receptor response spectra based on the hypothesis that the binding pocket properties partially encode this information, as suggested by our results on *Spodoptera littoralis* ORs.

## Introduction

Various insect behaviours are intimately linked to their sense of smell at all stages of their lives. In complex and ever-changing environments, insects must efficiently filter and interpret essential cues to locate food sources, potential mates, predators or suitable egg-laying sites^1^. These chemical cues are detected by seven transmembrane domain (TM) odorant receptor (OR) proteins located in the membrane of olfactory sensory neurons (OSNs) within olfactory organs. Unlike the mammalian ORs, which belong to the G protein-coupled receptor (GPCR) superfamily of seven TM proteins, insect ORs are odorant-gated ion channels directly responsible for signal transduction. They assemble in tetrameric complexes composed of two subunits: a conserved co-receptor (Orco) subunit and a highly divergent OR subunit^2,3,4,5,6^. The OR subunit, which houses the odorant-binding site, confers chemical selectivity to the heteromeric complex while Orco ensures subcellular OR trafficking, complex assembly, and participates in the pore architecture together with the variable OR^2,5,6,7,8,9^. Such an unusual ion channel structure has been confirmed by recent experimental structures obtained using cryo-electron microscopy (cryo-EM) ^2,3,5,6^.

Over the past two decades, the sequencing of a large number of insect transcriptomes and genomes has provided a wealth of OR sequences^10^. The number of ORs genes varies significantly among species, ranging from as few as 4 in damselflies^11^ to several hundred in social insects^12^. These genes evolve rapidly through a dynamic process of duplication, divergence, pseudogenization, and loss. This “birth-and-death” process results in the expansion or reduction of the OR repertoire reflecting the adaptation of insects to a diverse range of ecological niches^13^. Although ligands have now been identified for a large list of insect ORs using various heterologous expression systems^10^, ligand identification of many more ORs is needed to understand how a given species mobilizes its OR repertoire to interact with its environment. This will enhance our understanding of the molecular evolution of these ecologically important insect receptors.

Ligand-Based Virtual Screening (LBVS) represents a promising way to accelerate this process, by screening large databases and identifying the physico-chemical properties of an odorant required for activating a given OR^14,15,16,17,18,19^. Computational models can rapidly screen extensive libraries of relevant molecules, and the identified hits can move on to *in vitro* or *in vivo* testing for further validation. Traditional insect OR functional analyses typically limit the selection of odorants tested to those found in the animal’s environment^20,21^, potentially missing certain agonists. Integrating advanced computational methods into this process not only accelerates the discovery of effective molecules, but also expands the pool of potential candidates. The main limitation of LBVS is its reliance on ligand knowledge, which makes it applicable only to receptors that have already been deorphanized (i.e. ligand identified). However, the recent advances in our understanding of insect OR structures^2,3,5,6^, combined with the cutting-edge technology of AlphaFold^22^, provide an opportunity to implement Structure-Based Virtual Screenings (SBVS). Relying on docking simulations, SBVS has the potential to overcome LBVS barriers and greatly expand our ability to identify OR-odorant interactions *de novo*. SBVS is well established for mammalian GPCRs^23^, however its application towards insect ORs, which are unrelated to GPCRs, has yet to be demonstrated. This is primarily due to significant disparities in binding site structures. By November 2023, cryo-EM had resolved the structures of 523 GPCRs^24^, compared to only 5 insect ORs by 2024^2,3,5,6^. Currently, docking simulations on insect ORs have predominantly been employed to enhance our comprehension of the molecular mechanisms underlying olfactory signal transduction across various species by pinpointing receptor residues that may interact with known ligands, thereby providing hypotheses tested through targeted mutagenesis^3,5,6,25,26,27,28,29^. While few studies have begun to suggest the benefits of docking for predicting new ligands for ORs, none of them have experimentally validated their predictions^30,31^. To date, such complete investigations in insects have only been conducted on odorant-binding proteins (OBPs)^32,33,34^, a family of soluble proteins proposed to transport odorants to the OR^35^.

Here, we present a comprehensive study of virtual screening for insect ORs, spanning from *in silico* predictions to *in vivo* validations and behavioral assays. We screened a large database containing 407,270 natural compounds against two deorphaned ORs from the leafworm moth *Spodoptera littoralis* (Slit) with contrasting tuning breadths: SlitOR25, which exhibits a broad recognition spectrum, and SlitOR31, specifically tuned to interact with eugenol^20^. Recognized as a cornerstone model in chemical ecology, *S. littoralis* has been instrumental in providing valuable insights in this field for several decades. Thus, many odorants from this species’ host plants are known. Nevertheless, during our analysis we avoided making any assumptions about the chemicals responsible for activating these receptors, to increase our chance of discovering new, unexpected ligands. As a result, our SBVS approach led us to discover new OR ligands, one of which was localized in a previously unexplored area of the chemical space - an area that would remain inaccessible using LBVS methods. SBVS predictions were not only experimentally verified at the OR level, but their behavioral relevance was also investigated, pinpointing new insect attractants. We also highlight the need for meticulous evaluation of raw output generated by docking software since such data often fails to distinguish directly between agonists and non-agonists. Ultimately, the results of our predictive workflow, demonstrated across both broadly and narrowly tuned receptors, pave the way for future studies to cover any OR within *S. littoralis* and we anticipate applying this methodology to ORs of additional insect species.

## Methods

### Chemoinformatics and preparation of chemical libraries

The experimental dataset used to assess the docking protocol comprised 51 volatiles tested on various SlitORs in de Fouchier *et al*., 2017^20^ (Supplementary Table 1). The Structure-Based Virtual Screening was conducted using the COCONUT online database^36^. This open collection of natural products contains 407,270 molecules (latest updates: January 2022). We filtered the molecules according to the following criteria: retaining those with a molecular weight not exceeding 400 g/mol, featuring no more than 25 heavy atoms, having no more than 10 heteroatoms, and with a calculated octanol/water partition coefficient (logP) ranging between -1 and 7. These physico-chemical properties can be useful for distinguishing between odorous and odorless molecules, as observed in previous databases^37^. These molecular descriptors were calculated for all the compounds in the database using the RDKit python library (version 2023.03.3)^38^. Ultimately, 125,620 molecules were retained for the virtual screening (Fig. 1).

**Figure 1:**
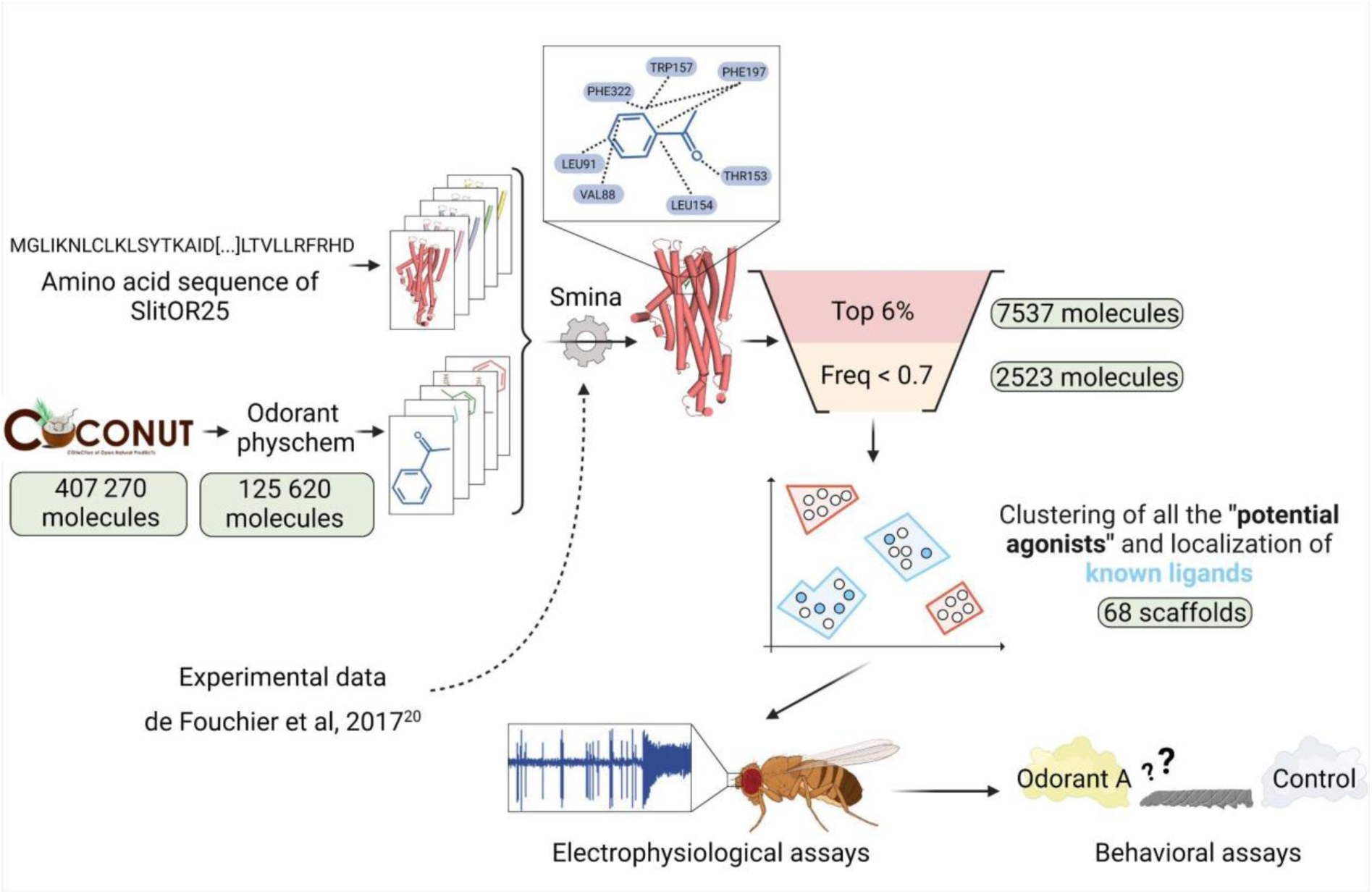
Structure-Based Virtual Screening workflow illustrated with SlitOR25. Starting from a large database of 407,270 natural compounds, various filtering and molecular docking steps led to a selection of less than 100 molecules for experimental validation using electrophysiological assays on the OR and behavioral assays on the larvae. Pale green labels indicate the number of molecules remaining after each step. *Odorant physchem* consists of filtering the COCONUT database^36^ with general physicochemical properties of volatile molecules (molecular weight, number of heavy atoms, number of heteroatoms, logP)^37^. “*Top 6%*” is a selection of the top 6 percent of docked compounds that exhibit the best docking scores. “*Freq < 0.7*” excludes from the selection the compounds that were in the top 6 percent for more than 70 percent of the SlitORs. The clustering was performed using HDBSCAN^57^. Scaffolds containing known ligands are highlighted in blue, while those defining unexplored regions of the chemical space are marked in red. Refer to the methodology section for more details.

Both sets of molecules -the one from de Fouchier *et al.*^20^ and the one from COCONUT^36^- were prepared using identical procedures. Gypsum-DL 1.2.0 was employed to generate 3D molecular structures from SMILES format to SDF^39^. All potential ionization, tautomeric, chiral, cis/trans isomeric, and ring-conformational forms were checked and optimized at a pH of 7.0 ± 0.5 using Gypsum-DL^39^. The conversion from SDF to MOL2 format was performed using Open Babel 3.1.0^40^, and the structures were subsequently processed from MOL2 to PDBQT using MGLTools (version 1.57)^41^.

### Modeling and analysis of SlitOR structures

Five relaxed models were generated with AlphaFold2^22^ for the 17 receptors of *Spodoptera littoralis* studied in de Fouchier *et al*., 2017^20^ SlitOR3, SlitOR4, SlitOR6, SlitOR7, SlitOR13, SlitOR14, SlitOR17, SlitOR19, SlitOR24, SlitOR25, SlitOR27, SlitOR28, SlitOR29, SlitOR31, SlitOR32, SlitOR35 and SlitOR36. From the models generated for each receptor, the one with the highest pLDDT score was chosen for docking studies. Polar hydrogen atoms were added using PDB2PQR 3.6.1^42^, with protonation states set by PROPKA 1.0^43^ at pH 7.0. The final structure was then minimized using the AMBER99 force field. MGLTools 1.5.7 were used to convert protein file format from PQR to PDBQT. Fpocket 4.0^44^ was employed for cavity detection, and MDpocket^45^ for the analysis of their physico-chemical properties. The structure of the ligand-receptor complexes were analyzed with ProLIF 2.0.3^46^. The reported findings were collected prior to the release of the latest AlphaFold version^47^. For comparative purposes, we have also generated models with AlphaFold3.

### Docking simulations

Cryo-EM structures of *Machilis hrabei* (Mhra) OR5 in complex with eugenol and DEET (Protein Data Bank accession 7LID and 7LIG) were superimposed with SlitOR models using PyMOL 2.5.4^48^. After visual inspection of the cavities generated by Fpocket^44^, the primary cavity coinciding with the MhraOR5 binding site was identified as the putative binding site for each SlitOR (Supplementary Fig. 1c, d). Residues within 5 Å from the center of these cavities were considered part of the binding sites. The grid box used for docking simulations was then optimized to fit with the potential binding sites of SlitORs, ensuring thorough coverage of cavity residues while minimizing their overall volume (center_x = 10, center_y =-1.67 center_z =-12, size_x = 13, size_y = 13.9, size_z = 15).

All the compounds were docked using the Vinardo scoring function implemented in the smina software^49,50^ (Oct 15 2019 based on AutoDock Vina 1.1.2) with an exhaustiveness value of 20, selected for its balance between speed and high performance^51^. The entropic penalty is sometimes neglected by the docking scoring function or poorly estimated based on the number of rotamers in the molecule^52^. Consequently, molecules with greater molecular weights tend to better occupy the binding pocket, forming more contacts with the protein, leading to increased docking scores. To address this issue, one approach involves weighing the score based on molecular weight^53,54^, similar to how ligand efficiency is calculated (by dividing the docking score by the number of heavy atoms)^55^. We re-evaluated Vinardo scores by weighting them in three different ways based on the number of heavy atoms in each molecule:

- LE score function: Vinardo Score Function / Number of Heavy atoms
- LEln score function: Vinardo Score Function / (1+ln(Number of Heavy atoms))
- LESA score function: Vinardo Score Function / (Number of Heavy atoms)^⅔^

ROC curves were generated for the smina outputs to evaluate the effectiveness of the score functions in distinguishing between agonists and non-agonists. The choice of the scoring function was determined by evaluating the docking performance, utilizing experimental data on the 17 modeled SlitORs (Fig. 1). To validate our choice, we performed an additional check by calculating the enrichment factor for all the scoring functions tested (Supplementary Table 2).

### Refining docking results and classification of predictions

To propose a suitable number of novel candidate ligands for testing on SlitOR25 and SlitOR31 in *in vivo* experiments, several filters were applied to select the most relevant compounds (Fig. 1). Leveraging experimental data gathered from the evaluation of 51 molecules by de Fouchier *et al*., 2017^20^, the true positive rate (TPR) to false positive rate (FPR) ratio was calculated for 15 SlitORs, namely SlitOR3, SlitOR4, SlitOR7, SlitOR14, SlitOR17, SlitOR19, SlitOR24, SlitOR25, SlitOR27, SlitOR28, SlitOR29, SlitOR31, SlitOR32, SlitOR35 and SlitOR36. We excluded from this analysis the pheromone receptors SlitOR6 and SlitOR13, as they did not require the same score function as the non-pheromonal receptors to achieve good discrimination between agonists and non-agonists (Fig. 1). By varying the threshold of the top x% of molecules retained, as ranked by their docking scores according to the LE function, the optimal TPR/FPR ratio was achieved when the threshold was set at 6%, which was the median optimal threshold value across all receptors. This threshold optimizes the differential categorization between agonists and non-agonists (Fig. 1) and was confirmed by analyzing the enrichment factors associated with the selected scoring function (Supplementary Table 2).

As a secondary filter, we excluded molecules showing remarkably strong interaction energy with almost every receptor, postulating that such promiscuity might result from a discrepancy in computational scoring rather than reflecting a biological reality. We analyzed the data from de Fouchier *et al.*, 2017^20^and calculated the occurrence frequency of molecules in the top 6% ranked with their LE scores. If a molecule appeared in the list for 70% of receptors, we found that it was likely to be a non-agonist. Therefore, we set the threshold at 0.7 (i.e. molecules predicted as agonists for 70% of ORs) and refer to the molecules discarded during this filtration step as “suspected decoys”.

Only compounds that successfully passed both filtration steps were considered “potential agonists” for the receptors tested. In re-examining de Fouchier’s experimental data^20^, we also calculated the median rank at which the True Positive Rate (TPR) equals 1 for the 15 non-pheromonal SlitORs. This indicates a threshold below which all molecules are predicted to be non-agonists. Consequently, we categorized molecules that fell within the lowest 45% of the docking score distribution as “potential non-agonists”.

### Clustering

To thoroughly explore the extensive chemical space and maximize the likelihood of discovering novel ligands with the most diverse structures possible, “potential agonists”, “potential non-agonists” and “suspected decoys” identified through virtual screening of the COCONUT database^36^ underwent separate clustering based on their molecular structures, following the same protocol. The Morgan2 fingerprints (2048-bit vector) of each molecule in the three sets were generated using the RDKit library^37^. Then, UMAP 0.5.4^56^ was employed to reduce the dimensionality down to 2 with parameters set to n_neighbors=10, min_dist=0.0, and default settings. This was followed by HDBSCAN 0.8.38^57^ for data clustering with a minimum cluster size of 30 and default parameters. The consistency of intra- and inter-cluster distances was evaluated by comparing the mean Tanimoto coefficient of molecules within the same cluster to the Tanimoto coefficient between the centroids of different clusters.

### Chemicals

The availability of molecules with the best docking scores in each cluster of “potential agonists”, “potential non-agonists” and “suspected decoys” was verified by querying the AMBINTER supplier database. Exceptionally, for clusters of “potential agonists” which contain already known agonists of SlitOR25 and SlitOR31, the availability of the top 5 molecules with the best docking scores was also checked. Selected compounds were purchased from AMBINTER (Orléans, France) and are listed in Supplementary Table 1. Depending on their solubility, odorants were either directly diluted in paraffin oil or passed through a stock solution in DMSO. We ensured that final dilution did not contain more than 5 % DMSO.

### Heterologous expression of SlitOR25 and SlitOR31 in *Drosophila melanogaster* olfactory neurons

Flies were reared on a standard cornmeal-yeast-agar medium and kept in Sanyo incubators (MIR-553) at a temperature of 25°C, under a 12-hour:12-hour light-dark photoperiod. The flies were transferred to 29°C for 24 hours before single-sensillum recording to optimize GAL4 activity while minimizing any impact on line viability^58^. The UAS-SlitOR25 and UAS-SlitOR31 fly lines were previously generated^20^. These two UAS lines were crossed with *Df(2L)Or22ab*, *TI{GAL4}Or22ab* stock flies to express the OR of interest in ab3 *Drosophila melanogaster* (Dmel) sensilla instead of OR22a^59^.

### Single-sensillum recordings of ab3 *Drosophila* sensilla

Male and female flies, aged 2 to 5 days, were randomly selected from the *Drosophila* population. The flies were immobilized inside a 200 µl pipette tip leaving only the head exposed. The pipette tip was affixed with dental wax onto a microscope glass slide, ensuring that the ventral side of the fly faced upwards. The antennae were secured in place using a glass capillary positioned between the second and the third antennal segment. The entire assembly was positioned under a microscope (BX51WI, Olympus, Tokyo, Japan) equipped with an SLMPLN 100X objective. A continuous flow of charcoal-filtered and humidified air at a rate of 1.5 L.min^−1^ was directed through a glass tube with a 7 mm diameter to prevent desiccation of the flies.

The stimulation cartridges used for the screening of the odorant panel were made by placing a 1 cm² filter paper loaded with 10 µl of each odorant solution (10 µg/µl) into a Pasteur pipette. To prevent unintended evaporation of volatile compounds, a 1000 µl pipette plastic tip was used to securely seal the Pasteur pipette. Odorant stimulations were performed by inserting the Pasteur pipette into an opening in the glass tube from which the continuous airflow was delivered, and initiating a 500 ms air pulse at a rate of 0.6 L/min.

The firing activity of ab3A OSNs was recorded using tungsten electrodes. The reference electrode was inserted into the fly’s eye using a manually controlled micromanipulator (Three-Axis Coarse Mechanical Micromanipulator UM-3C, Narishige, Tokyo, Japan). The recording electrode was inserted at the base of the recorded sensillum using a motor-controlled micromanipulator (PatchStar Micromanipulator, Scientifica, Uckfield, United Kingdom). The electrical signal was amplified using an EX-1 amplifier (Dagan, Minneapolis, Minnesota, United States of America), digitized through a Axon Digidata 1550A Data Acquisition System (Molecular Devices, Sunnyvale, California, United States of America), recorded and analyzed using the pCLAMP 10.5 software (Molecular Devices). Net responses of ab3A neurons expressing SlitOR25 or SlitOR31 were determined by subtracting the spontaneous firing rate from the firing rate during the odorant stimulation, following the methodology outlined in de Fouchier et al., 2017^20^. To distinguish ab3 from other basiconic sensilla, 3 diagnostic stimuli were employed: 10 µg of sulcatone, 10 µg of ethyl acetate and CO_2_ delivered by human expiration. These stimuli are potent activators of ab3B OSNs, ab2A OSNs and ab1C OSNs, respectively^60^. The absence of the endogenous receptor OR22a in the ab3A neuron was confirmed by employing 10 µg of ethyl hexanoate, a strong ligand of DmelOR22a. Additionally, the presence of the SlitORs in the transformed ab3A OSNs was confirmed using known ligands as positive control: acetophenone for SlitOR25 and eugenol for SlitOR31^20^.

For each of the two receptors, the entire panel was tested across 7 different flies. A compound was deemed active if the neuronal response it elicited was statistically different from the response induced by the solvent alone (Friedman test, followed by a Dunnett’s multiple comparison test adjusted with Benjamini & Hochberg correction, p < 0.05). Dose-response experiments were conducted for all identified ligands, ranging from 1 ng to 100 µg. Data were recorded and analyzed using pCLAMP 10 and all statistical analyses were performed using R (version 4.3.2).

### Larvae behavioral assays

The potential attractiveness of the new compounds that activated SlitOR25 during the electrophysiological assays were tested using Y-tube olfactometers. We did not test the new ligand of SlitOR31 because eugenol, the primary ligand for this receptor, is known to be non-attractive to the larvae of *S. littoralis*^61^.

Two-choice behavioral assays were performed at 24°C under dim red light using 12-15 day old larvae (third or fourth instar, L3-L4) reared on artificial diet. Caterpillars were starved the night before the experiments (16 to 20 hours starvation). It was shown that this condition did not impact the survival or mobility of the caterpillars but increased the interest of the caterpillars in odor sources^62^. The olfactometer consisted of a glass Y-tube with an internal diameter of 2.1 cm. The main segment was 13 cm long and each of the two arms was 9.5 cm long. The air passing through the system was purified with charcoal. A flow meter (ProFLOW 6000, Restek, Bellefonte Pennsylvania, USA) ensured that the flow rate in each arm of the olfactometer was kept constant (0.5 L/min). Each of the tested odorants was diluted in paraffin oil at 10, 1 and 0.1 µg/µl. 10 µl of these solutions were loaded on a filter paper in one of the two arms of the Y-tube. 10 µl of paraffin oil was deposited on the filter paper in the other arm of the olfactometer. As the attractiveness of benzyl alcohol has already been demonstrated^61^, 100 µg of this compound was used as a positive control. Neutral controls consisted of paraffin oil in both arms, and paraffin oil versus (E)-ocimene, a compound that does not induce any *S. littoralis* larval behavior^61^.

Each caterpillar was tested only once, each olfactometer was changed every 4 caterpillars, and the two arms of the olfactometer were interchanged between individuals. After each experiment, all the glass parts of the device were cleaned for 1 hour in a 3-5 % solution of detergent TDF4 (Franklab, Montigny-le-Bretonneux, France), rinsed with distilled water, dried, rinsed with acetone, and put in an oven at 200 °C for 1 hour.

Pearson’s Chi-squared tests were used to confirm whether the proportion of larvae preferentially choosing one arm of the olfactometer differed from 50:50 (p<0.05). The choice of a caterpillar was recorded if three quarters of its body was visible in one of the arms of the olfactometer within10 minutes.

## Results

### Selecting the appropriate scoring function enhances docking performance in discriminating odorant receptor agonists and non-agonists

To determine the most effective scoring function for distinguishing potential agonists from non-agonists in the COCONUT database^36^, we carried out a preliminary comparison. The aim was to assess the performance of different scoring functions using experimental data from de Fouchier *et al*., 2017^20^, wherein the responses of 17 *S. littoralis* ORs to 51 odorants, including plant volatiles and sex pheromone compounds, were measured. We plotted receiver operating characteristic (ROC) curves and calculated the area under the curve (AUC) for each receptor^63^. First, we separated our analysis between pheromone receptors (PRs, tuned to sex pheromone compounds) and non-pheromonal receptors due to their divergent responses (Fig. 2a). Notably, the AUC values of ROC curves were markedly high for PRs when utilizing the Vinardo scoring function, with a median value of 0.90. Conversely, this value was considerably lower for non-pheromonal ORs, with a median value of 0.39. Consequently, the Vinardo scoring function could not efficiently discriminate between agonists and non-agonists for non-pheromonal receptors like SlitOR25 and SlitOR31. Weighting the Vinardo scoring function by taking into account the number of heavy atoms (LE scoring function) resulted in a remarkable improvement for SlitOR25 and SlitOR31, almost doubling the AUC values to 0.87 and 0.72, respectively (Fig. 2a-b). Such a substantial improvement was observed for all non-pheromonal receptors, with a median AUC value of 0.72. The increase in the AUC values with the LESA and LEln scoring functions were less pronounced, reaching 0.48 and 0.65 for SlitOR25 and SlitOR31, respectively (Fig. 2a-b).

**Figure 2:**
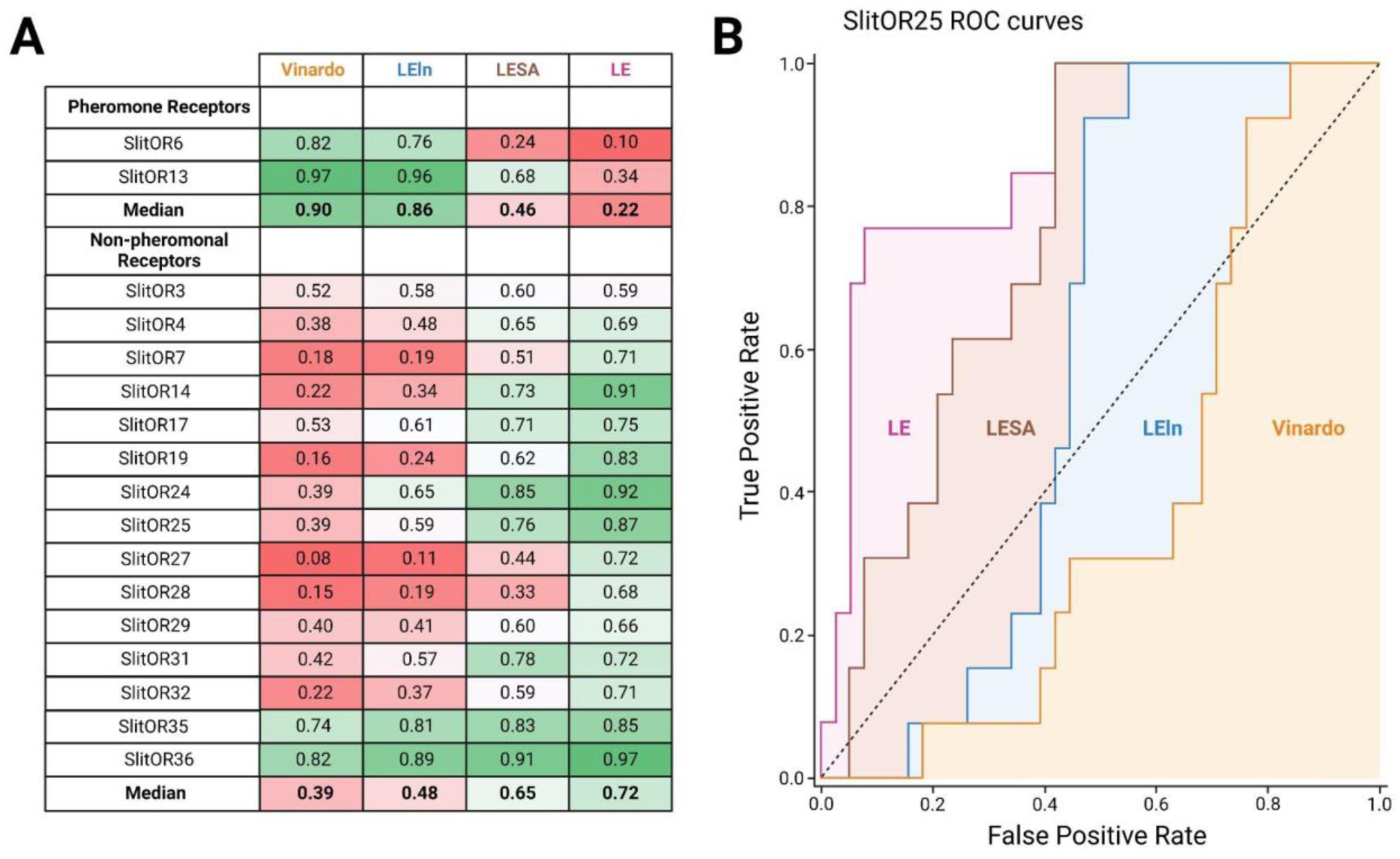
LE stands out as the top-performing scoring function for analyzing the odorant docking poses on non-pheromonal odorant receptors of *S. littoralis*, including SlitOR25 and SlitOR31. (**A**) The comparison table summarizes AUC values from ROC curves using Vinardo, LESA, LEln, and LE scoring functions based on optimal poses of 51 molecules across 17 receptors from de Fouchier et al., 2017^20^. The LESA, LEln, and LE functions are weighted versions of the Vinardo function as described in methodology. The gradient color scheme highlights AUC values, with green indicating higher performance and red lower ones. (**B**) ROC curves for SlitOR25 show performance with Vinardo (orange), LEln (blue), LESA (brown), and LE (pink) scoring functions.

Equivalent results were obtained from the calculation of enrichment factors associated with the different scoring functions evaluated in this study. The maximum enrichment factor values were high for PRs using the Vinardo scoring function, with a median value of 8.93, and high for other ORs using the LE scoring function, with a median value of 5.67. Notably, replacing the Vinardo scoring function with the LE scoring function resulted in a substantial improvement for SlitOR25 and SlitOR31, increasing their maximum enrichment factors by approximately threefold and sixfold, respectively (Supplementary Table 2).

During the screening of the COCONUT database, the LE scoring function was then employed to assess the docking poses of the 407,270 molecules.

### Virtual screening of a large database opens unexplored chemical spaces of novel active compounds

After implementing an optimal docking procedure, we narrowed down 407,270 natural compounds to 2,523 potential agonists for SlitOR25 and 2,516 for SlitOR31. These subsets consisted of the top 6% scoring compounds devoid of probable decoy molecules, i.e. removing those interacting with more than 70% of non-pheromonal receptors. The chemical space of SlitOR25 and SlitOR31 was depicted using UMAP^56^ based on the chemical structure (Morgan2 fingerprint) of potential agonists. HDBSCAN clustering^57^ grouped these molecules into 68 clusters for SlitOR25 and 70 clusters for SlitOR31 (Supplementary Fig. 2). Clusters containing natural compounds with previously known experimental activity on SlitOR25 or SlitOR31 have been identified (Fig. 3). Molecules with top docking scores were chosen to represent each cluster. For experimental assays, we prioritized those purchasable and localized in the most densely populated clusters (Fig. 3). The 19 potential agonists selected for testing on Slit*OR25* fell into 15 clusters. Eight molecules were localized within four clusters housing known agonists, while 11 belonged to clusters never explored for SlitOR25. Among the 17 molecules picked for SlitOR31 testing, four were within the cluster housing the sole known agonist for this receptor, while the remaining ones were spread across 13 unexplored clusters. For quality-control purposes, we also selected 3 suspected decoys and 5 potential non-agonists for *in vivo* testing to validate that the various filtration steps did not eliminate ligands of SlitOR25 and SlitOR31. They were chosen following a procedure similar to the one described above. Within each cluster of suspected decoys and potential non-agonists, the molecule with the best docking score was retained. From this selection, decisions were based on supplier availability, with a focus on molecules within largest clusters.

**Figure 3:**
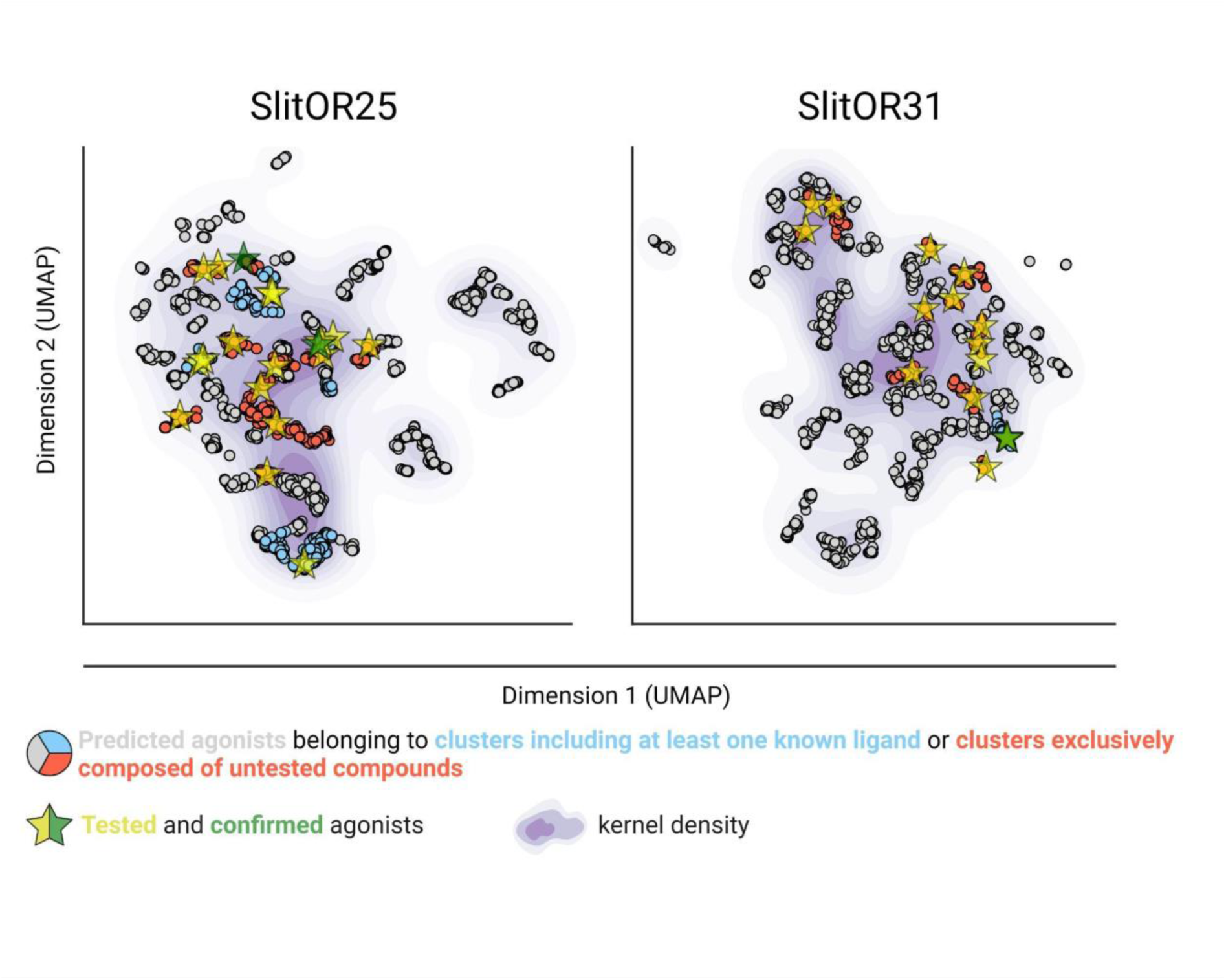
Virtual screening allowed the identification of new ligands in unexplored areas within the predicted chemical space of SlitOR25 and SlitOR31. UMAP representation of the chemical space based on Morgan2 fingerprints of potential agonists, and with kernel density estimates illustrated using a purple gradient. All the potential agonists experimentally tested are represented by yellow stars. These stars are recolored in green when the tested compound is confirmed to be a real ligand for the receptor of interest.

### *In vivo* experiments confirm the identification of new ligands for both the generalist SlitOR25 and the specialist SlitOR31 receptors

To validate the *in silico* predictions, we conducted single-sensillum recordings on *Drosophila* ab3A OSNs expressing SlitOR25 or SlitOR31 in place of the endogenous DmelOR upon stimulation with the predicted agonists, decoys and non-agonists.

SlitOR25 exhibited significant activation in response to 2 out of the 19 molecules predicted as agonists by docking (Fig. 4a): benzyl formate (labeled 16 in Fig. 4a and in Supplementary Table 1) and 3-(3,4- dihydroxyphenyl)propanoic acid (10). This corresponded to a success rate (precision) of over 10%. Furthermore, with an accuracy of 0.36, the virtual screening campaign has effectively ruled out a significant number of potential non-active compounds, as none of the selected molecules predicted to be non-agonists or suspected decoys activated SlitOR25. Additional metrics evaluating the performance of the workflow are summarized in Supplementary Table 3. It should be noted that the median response of neurons expressing SlitOR25 to benzyl formate, at 162 spikes/s, was comparable to the response elicited by acetophenone^20^ or by benzyl cyanide^16^, the best SlitOR25 ligands previously identified. Benzyl formate (16) belonged to a cluster of molecules containing SlitOR25 previously identified ligands^16,20^. Remarkably, 3-(3,4-dihydroxyphenyl)propanoic acid (10) was part of a new cluster (Fig. 3 and Supplementary Fig. 2). By comparing this new ligand with known agonists, we found that the maximum Tanimoto coefficient obtained is low (Tc=0.25), revealing unusual chemical structures for a SlitOR25 ligand. Dose-response experiments were conducted for the two new SlitOR25 ligands discovered in this study (Fig. 4b-c). Compound 10 showed significant receptor activation only at 100 µg. SlitOR25 appeared to be more sensitive to compounds 16, which significantly activated the receptor starting from 10 µg.

**Figure 4:**
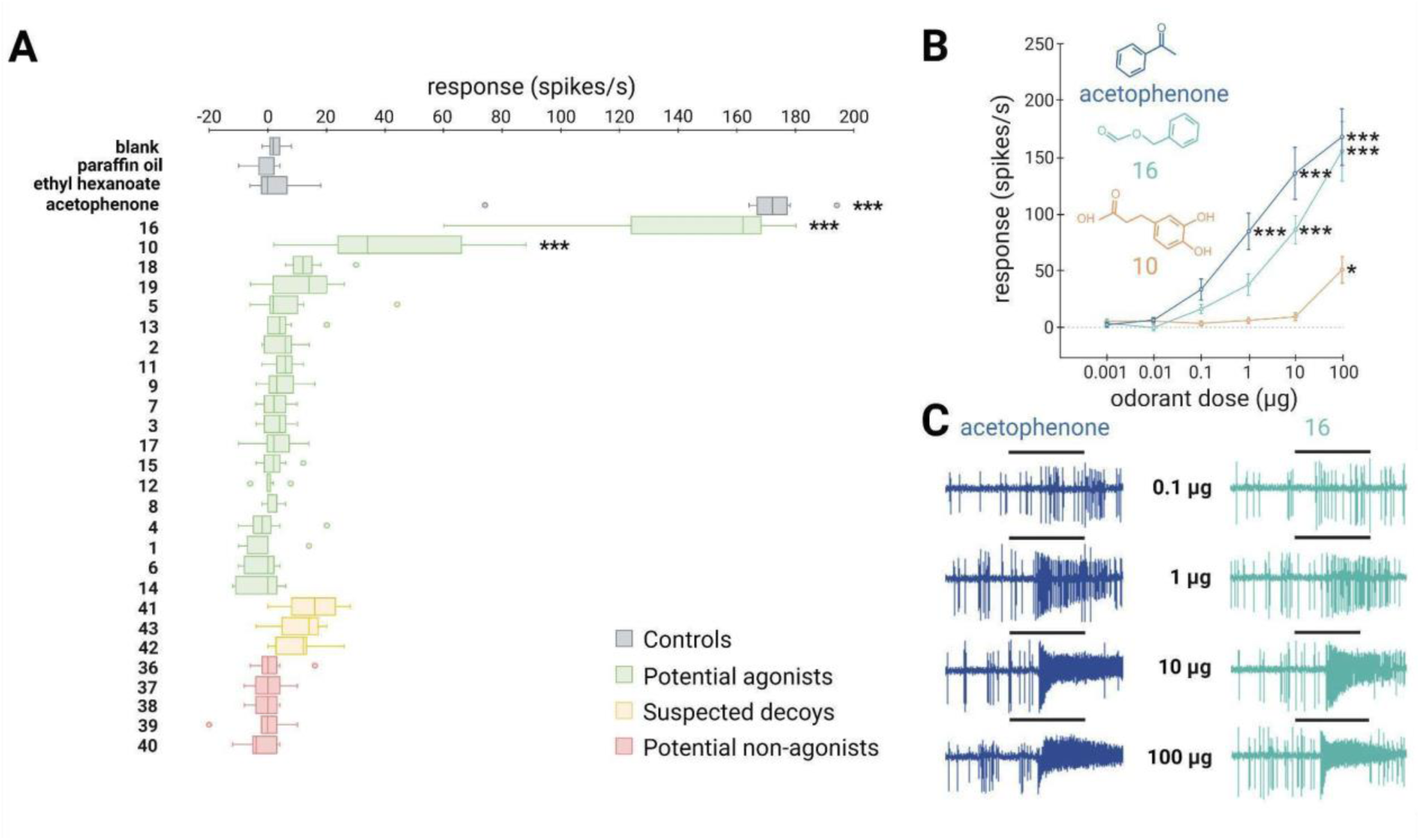
∼10% of the predicted agonists via the SBVS approach activate the broadly-tuned receptor SlitOR25. (**A**) Box plot shows the responses of *Drosophila* ab3A OSNs (n = 7) expressing SlitOR25 measured upon exposure to a panel of predicted agonists, decoys, and non-agonists (100 µg loaded in the stimulus cartridge). The whiskers were extended to 1.5 times the interquartile range from the quartiles. Outliers are indicated by dots. The controls are represented in grey (blank, paraffin oil solvent, ethyl hexanoate and acetophenone). Green boxes represent the responses to predicted agonist compounds, yellow boxes depict the responses to predicted agonist compounds that have been removed from our selection because they activate the majority of the tested SlitORs (suspected decoys), and red boxes indicate the responses to predicted non-agonist compounds. (**B**) Dose-response curves of *Drosophila* ab3A OSNs for all active compounds revealed during the screening of the panel. Data represented are mean action potential frequencies ± SEM (*n* = 7). In (**A**) and (**B**), asterisks indicate statistically significant differences between responses to the odorant and to the solvent (Friedman test followed by a Dunnett’s multiple comparison test with a Benjamini & Hochberg correction; \**p* < 0.05, \*\**p* < 0.01, \*\*\**p* < 0.001). (**C**) Examples of single-sensillum recording traces obtained for *Drosophila* ab3A OSNs expressing SlitOR25 stimulated with increasing doses of acetophenone (known ligand) and benzyl formate (16, new active compound which exhibited the highest response). Black bars represent the duration of the stimulus (500 ms).

Out of the 17 molecules predicted to activate SlitOR31, only 2-methoxy-4-propylphenol (23) induced significant responses, resulting in a success rate of approximately 6% (Fig. 5a). Compounds 23 was part of a cluster that also contained eugenol, the only known ligand of SlitOR31 (Fig. 3 and Supplementary Fig. 2). In line with the findings for SlitOR25, no significant response was observed for the selected molecules predicted as non-agonists or suspected decoys. To delve deeper into SlitOR31 sensitivity to compound 23, we conducted dose-response experiments (Fig. 5b-c). This compound significantly activated SlitOR31, with activity observed starting from 10 µg. Eugenol remained the most potent SlitOR31 ligand, initiating receptor activation at 1 µg.

**Figure 5:**
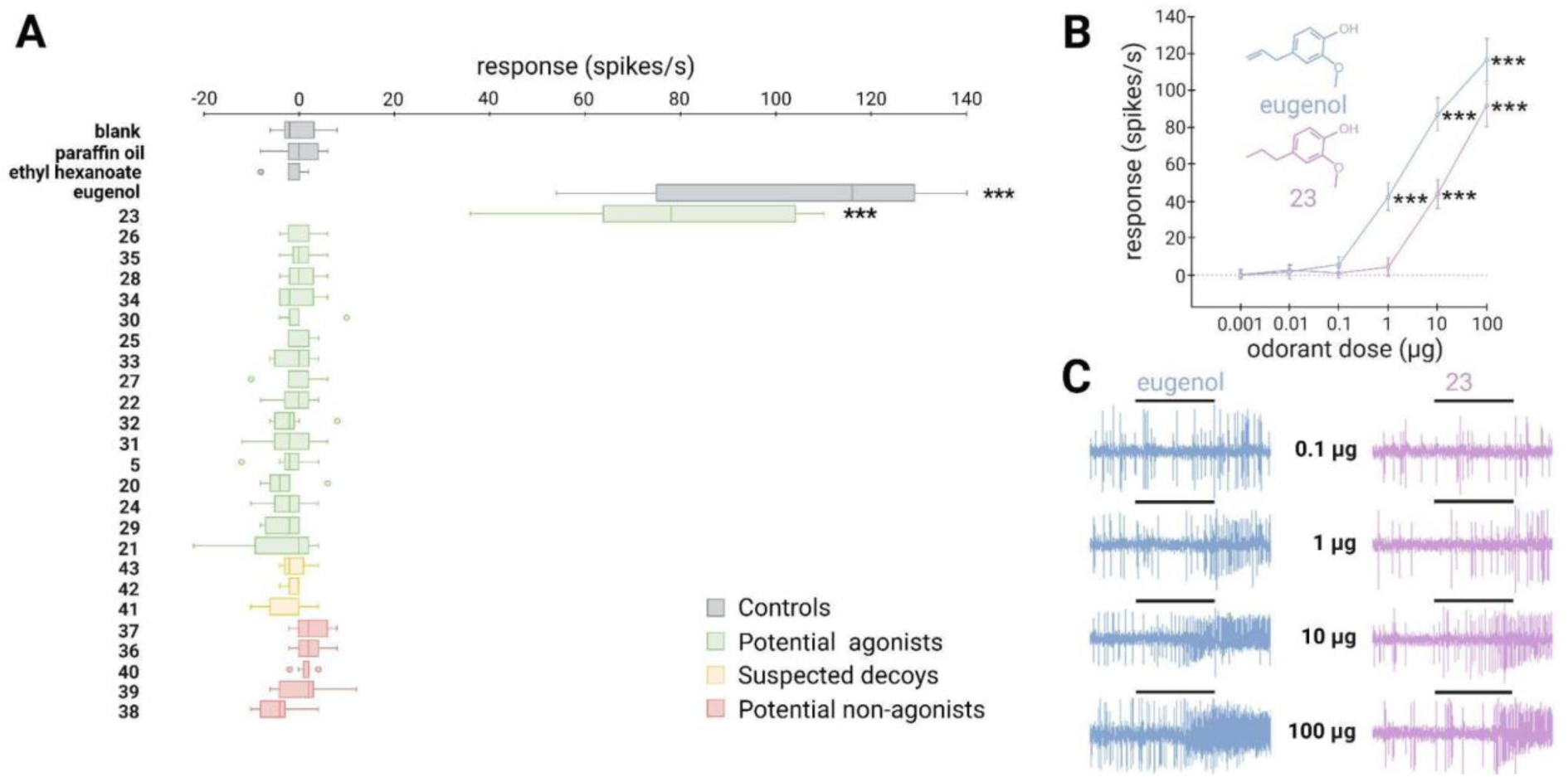
A new agonist predicted via the SBVS approach activates the narrowly-tuned receptor SlitOR31. (**A**) Box plot shows the responses of *Drosophila* ab3A OSNs (n = 7) expressing SlitOR31 measured upon exposure to a panel of predicted agonists, decoys, and non-agonists (100 µg loaded in the stimulus cartridge). The whiskers were extended to 1.5 times the interquartile range from the quartiles. Outliers are indicated by dots. The controls are represented in grey (blank, paraffin oil solvent, ethyl hexanoate and eugenol). Green boxes represent the responses to predicted agonist compounds, yellow boxes depict the responses to predicted agonist compounds that have been removed from our selection because they activate the majority of the tested SlitORs, and red boxes indicate the responses to predicted non-agonist compounds. (**B**) Dose-response curves of *Drosophila* ab3A OSNs for all active compounds revealed during the screening of the panel. Data represented are mean action potential frequencies ± SEM (*n* = 7). In (**A**) and (**B**), asterisks indicate statistically significant differences between responses to the odorant and to the solvent (Friedman test test followed by a Dunnett’s multiple comparison test with a Benjamini & Hochberg correction; \**p* < 0.05, \*\**p* < 0.01, \*\*\**p* < 0.001). (**C**) Example of single-sensillum recording traces obtained for *Drosophila* ab3A OSNs stimulated with increasing doses of eugenol (known ligand) and 2- methoxy-4-propylphenol (23, new active compound). Black bars represent the duration of the stimulus (500 ms).

### The binding sites of SlitOR25 and SlitOR31 differ both in their amino acid composition and interaction patterns

To further understand why the broadly tuned receptor SlitOR25 and the specific receptor SlitOR31 exhibited divergent chemical spaces, we thoroughly explored the conservation of their binding site and the interactions of all compounds identified as agonists from this and previous studies^16,20^. Data from 33 molecules for SlitOR25 and two for SlitOR31 were compiled. While residues forming the pore of the ion channel in the OR/Orco complex were significantly conserved within the SlitORs, the residues forming the binding sites exhibited higher variability (Fig. 6a-c). These differences could potentially correlate with the diverse chemical spaces detected by the SlitORs.

**Figure 6:**
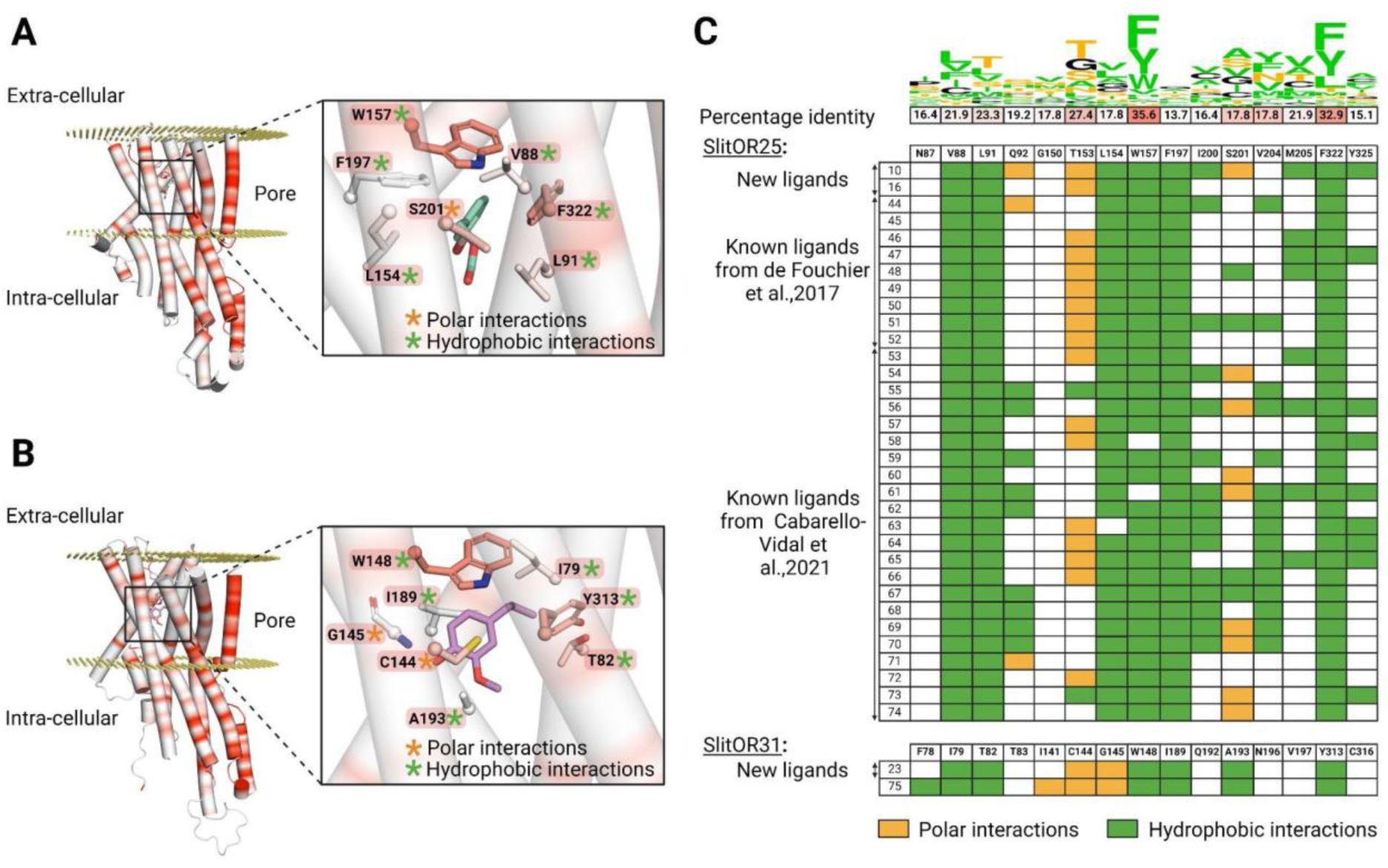
Activation patterns differ between a broadly-tuned receptor (SlitOR25) and a specific receptor (SlitOR31). (**A**) Structure of SlitOR25 and (**B**) SlitOR31 respectively interacting with benzyl formate (16) and 2- methoxy-4-propylphenol (23), the newly discovered ligands inducing the most robust responses from their respective receptor. The receptor orientation in the membrane was determined using OPM server^64^. Orange and green asterisks symbolize polar and hydrophobic interactions, respectively, formed between the ligand and seven residues within the receptor’s binding pocket. Additionally, a gradient color coding across the entire protein highlights the conservation of amino acids among the 73 odorant receptors of *S. littoralis*: highly conserved residues appearing in red and non-conserved ones in white. The backbones of the residues are shown only if they interact with the molecules; otherwise, only the side chains are visualized. This figure was generated using the molecular visualization software PyMol^48^. (**C**) Comprehensive overview of ligand-receptor interaction pattern across all known natural compounds activating SlitOR25 and SlitOR31. We utilize the same color code as used in (**A**) and (**B**): hydrophobic interactions are depicted in green and polar interactions in orange. At the top of this table, the percentage of identity among interacting residues within the 73 SlitORs is indicated and coupled with a Show Logo. Hydrophobic residues are highlighted in green. Residues that can potentially form polar interactions are highlighted in orange, except for Y and W that are already colored in green. The others are in black.

SlitOR25 ligands appeared to bind mainly via hydrophobic interactions, and following a non-conserved ligand-receptor interacting scheme. The following thirteen residues interacted with at least one ligand: V88, L91, Q92, T153, L154, W157, F197, I200, S201, V204, M205, F322, and Y325. As shown in Fig. 6a-c, a majority of active compounds were also engaged in a unique polar interaction mainly with T153 and/or S201. The new ligand (10) is a notable exception, exhibiting three hydrogen bond interactions. It is reasonable to classify these polar interactions as hydrogen bonds even if some do not fully meet all structural criteria for conventional hydrogen bonds and should be considered as weak hydrogen bonds. Interestingly, SlitOR25 activation appeared to be versatile, as it did not strictly adhere to a predefined interaction pattern. Six residues (V88, L91, L154, W157, F197, and F322) constituted the core of the binding site and exhibited hydrophobic interactions with over 90% of the known ligands. Among them, W157 and F322, which form the binding pocket’s lid, were the most highly conserved residues in the binding pocket across SlitORs (Fig. 6c). The aromatic amino acid W157 aligned with W158 in the MhraOR5 structure^3^ and with F115 in the *Acyrthosiphon pisum* (Apis) OR5 structure^5^ which are key residues for ligand detection. Moreover, residue W157 also aligned with F136 in the *Aedes aegypti* (Aaeg) OR10 structure^6^ where its rotation enables the transition between the inactive and active states of the receptor. Conversely, residues Q92, I200, S201, V204, M205 and Y325 displayed hydrophobic interactions with less than 46% of the ligands, suggesting that the binding pocket has the capacity to generate additional anchoring points for various odorant structures, in line with the receptor’s wide range of recognition capabilities.

Looking at SlitOR31, 10 residues (F78, I79, T82, I141, C144, G145, W148, I189, A193 and Y313) were found to interact with the ligands. The ligand binding was mediated by hydrophobic and polar interactions that were widely shared across known ligands (Fig. 6b, c). Some structural features of the binding pocket were common between SlitOR31 and SlitOR25. In each case, the ligands were stabilized through hydrophobic interactions with a subset of aliphatic and aromatic side chains. As for SlitOR25, W158 and Y313, which are among the most conserved residues across SlitOR binding pockets, were observed to interact with every ligand and contribute to form the binding pocket’s lid. Here again, polar interactions might involve either typical hydrogen bonds or weaker variations, such as the polarized S-H bond in the C144 thiol group, which exhibits reduced polarity compared to a standard hydrogen bond. In contrast, SlitOR25 residues interact via their side chains, while interactions with SlitOR31 residues I141 and G145 occurred through backbone atoms and polar functional groups of the odorants.

In total, ten residues of the SlitOR25 binding pocket and nine residues of the SlitOR31 binding pocket were aligned with amino acids in the binding pockets of MhraOR5^3^, ApisOR5^5^, AaegOR10^6^ or DmelOrco^65^, where mutations have been shown to significantly impact ligand-receptor interactions (Supplementary Table 4). Therefore, we hypothesize that these residues play a critical role in the activation of these two receptors.

### The two new ligands of SlitOR25 are highly attractive to *S. littoralis* larvae

The electrophysiological tests have confirmed or refuted the predicted activity of potential agonists identified by our SBVS workflow on SlitOR25 and SlitOR31. To go further, we next investigated the biological relevance of the newly identified ligands on *S. littoralis* behavior using a dual-choice assay in a Y-tube olfactometer. Since previous studies have demonstrated that SlitOR25 activation leads to larvae attraction^16,61^, we speculated that the new ligands would induce a similar behavior. By contrast, the primary ligand of SlitOR31, eugenol, being not known to induce any specific larvae behavior^61^, we did not investigate the effect of the new SlitOR31 ligand. The viability of our experimental protocol was verified using three control compounds: paraffin oil (solvent), benzyl alcohol (known attractant), and (E)- ocimene (known to be neutral). All controls resulted in the expected larval responses: no choice for a specific arm in paraffin oil vs paraffin oil and paraffin oil vs (E)-ocimene assays, choice to the odorized arm in paraffin oil vs benzyl alcohol assay (Fig. 7). We next tested the effects of the new SlitOR25 ligands, benzyl formate (16) and 3-(3,4-dihydroxyphenyl)propanoic acid (10) vs paraffin oil on larval choice, and at different doses : 100, 10, and 1 µg. At 100 µg, 76.7% and 71.4% of the larvae made a choice to the arm odorized with compounds 16 and 10, respectively. Compound 16 retained activity at 10 µg, with a choice percentage of 67.7, but not at 1 µg, whereas compound 10 did not induce any choice at 1 and 10 µg. These results correlate with the difference we observed in SlitOR25 sensitivity to the two compounds during the SSR recordings, where the detection threshold of SlitOR25 was lower for benzyl formate (16) than for 3-(3,4-dihydroxyphenyl)propanoic acid (10).

**Figure 7:**
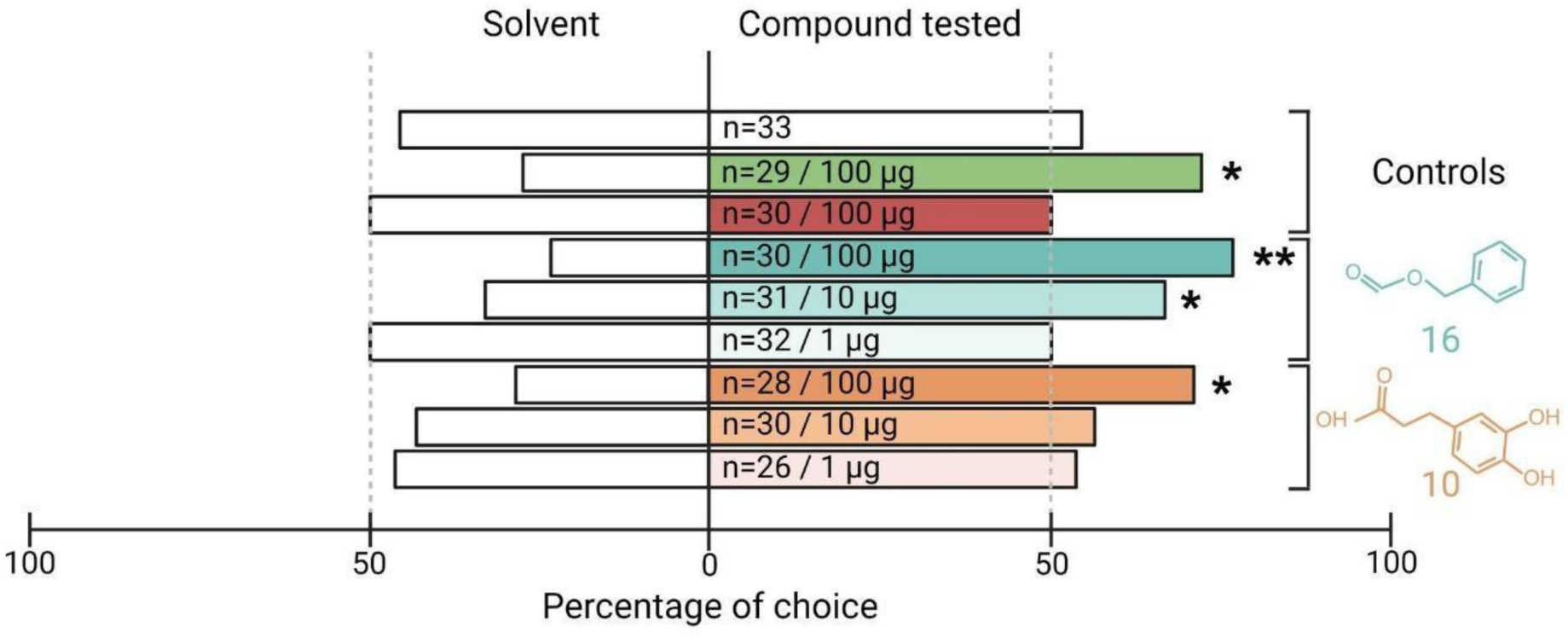
The two new ligands of SlitOR25 discovered through our Structure-Based Virtual Screening workflow are attractive for the larvae of *S. littoralis*. Data are presented as a percentage of larval choice. Controls: blank control (paraffin oil/paraffin oil, white bar), positive control (paraffin oil/benzyl alcohol, green bar), neutral control (paraffin oil/(E)-ocimene, red bar). The choices (paraffin oil/SlitOR25 new ligands) related to benzyl formate (16) and 3-(3,4-dihydroxyphenyl)propanoic acid (10) are represented in cyan and orange, respectively. The tested doses and the number of replicates (n) are indicated inside each bar of the diagram. Asterisks indicate statistically significant preferences of larvae for the odorized arm (Chi-squared test; *p < 0.05, **p < 0.01).

## Discussion

A better understanding of an animal chemical ecology requires extensive knowledge on the odorants it perceives and the resulting effects. Traditionally, chemical ecology approaches examine naturally occurring chemically-mediated processes in the species environment, focusing on known food diet, animal, and environmental volatiles ecologically relevant to the targeted species, following an *a priori* strategy. On the contrary, reverse chemical ecology investigates the underlying mechanisms to identify the proteins involved (e.g. odorant receptors), to be used as targets for large screens without any a priori on ecological relevance, offering the possibility to explore new areas of chemical spaces. In that view, investigating novel chemicals targeting insect ORs holds considerable promise for expanding our fundamental understanding of their chemical ecology and enabling practical advancements in pest control *via* the development of new behavioral disruptors. In the present work, we set up a joint computational and experimental approach to extend the chemical spaces of ORs from the crop pest moth *Spodoptera littoralis*. In particular, we focused on one broadly and one narrowly tuned OR to virtually screen a large database of natural compounds, coupled to experimental validation of the predicted ligands and behavioral assays. Unlike ligand-based methods which rely on prior knowledge of ligand structures, the Structure-Based Virtual Screening pipeline we propose here efficiently processed large libraries and utilized predictive modeling that incorporated established molecular recognition principles to discover compounds in both explored and unexplored chemical spaces. To our knowledge, this is the first report of a SBVS approach successfully employed for insect ORs. With a success rate ranging from 6% to 10% in accurately predicting active compounds, we demonstrate the power of the approach. As underlined in a recent review dedicated to docking simulations, a hit rate of 5% or even 3% is considered more than acceptable^66^. This arises when limited information about the target is available or when the screening library has not undergone any preliminary filtering. Higher performance can be reached if an experimental structure is available or, for instance, the location of the binding site or key interacting residues are known. Moreover, our approach not only identified potential agonists but also effectively distinguished non-agonists.

However, the SBVS approach is far from perfect and fails in some cases. It may be due to discrepancies in the AlphaFold2-predicted structure of the binding pocket, as has recently been highlighted^67,68^. Comparing the models we obtained using Alphafold2 with those obtained through the newly released version Alphafold3^47^, which has been specifically designed for addressing this issue, revealed no significant improvement. The RMSd between Alphafold2 and Alphafold3 models was always lower than 1.2 Å for the entire proteins and lower than 0.4 Å on the binding pocket (Supplementary Fig. 1a-b). It would have been possible to explore the gains associated with considering flexible receptor docking for the experimental dataset consisting of 51 molecules. However, this approach would have significantly increased both computation time and resource requirements for the database containing several hundred thousand molecules. Indeed, the current trend is not to complexify the docking algorithm but rather to apply it on a subset of a large library by prioritizing molecules that occupy the same chemical space as potential hits (by calculating chemical similarity and clustering)^69,70^ or by training a machine learning model capable of screening the entire database^71,72^. We can then consider that the scoring function alone is not sufficient to accurately rank and distinguish agonists and non-agonists. As shown in Fig. 2, the Vinardo scoring function performed well for pheromone receptors and poorly for non-pheromonal ORs, which led us to recalibrate the scoring function. It should be noted that the formation of a ligand-receptor complex is driven by the free energy of binding, which can be decomposed into two components: an enthalpic and an entropic contribution. In docking scoring functions, the entropic component may not be accurately accounted for or even overlooked entirely, often being approximated solely on the basis of the number of rotatable bonds within the molecule. As a result, large molecules with high molecular weights are typically able to fit better into the binding site and to establish more interactions with the target protein, resulting in higher docking scores. This is what we suspected by comparing docking scores with *in vivo* responses for various classes of ligands (Supplementary Fig. 3). A potential solution to the overestimation of the enthalpic term (or an underestimation of the entropic penalty) was to adjust the scoring function so that it takes into consideration the molecular weight (or the number of heavy atoms) of each compound. We tested various rescoring methods and observed that weighting, such as the Ligand Efficiency (LE) score, provides the best performance. It is not so usual to replace the docking scoring function by LE but it has already been useful in previous studies^53,54,55^. However, using Ligand Efficiency scoring reversed the tendency and wrongly ranked very small molecules as top ligands. This suggests that solving the problem would require the optimization of a new scoring function including a parameter accounting for molecular weight. Moreover, the rescoring did not have the same effect on all the receptors. For instance, when considering SlitOR3, the improvement was modest, as shown in Fig. 2a (AUC increases from 0.52 to 0.59 when using LE instead of Vinardo). A possible rationale lies in the application of a rigorous statistical criterion (*p*-value below 0.001) for categorizing compounds as either agonists or non-agonists. Adopting a less stringent *p*-value threshold enhanced docking performance, as demonstrated in Supplementary Fig. 4. This observation underscores the significance of carefully curating databases tailored for building meaningful computational models. So, to avoid a bias in the optimization function due to a limited dataset of 850 ligand-receptor pairs (51 odorants tested with 17 SlitORs), we finally decided to define a specific filter on the docking output results to exclude molecules displaying top rank position across nearly all receptors as it might actually be attributed to a scoring error, rather than depicting true biological activity. Defining such a “suspected decoy” filter effectively reduced the number of false positives and may be a viable compromise to maximize docking performance without the need to create a new scoring function from scratch.

Beyond its ability to explore new chemical spaces, the strength of the SBVS approach lies in its capability to unravel structure-function relationships, which sets it apart from LBVS. For instance, SBVS can guide site-directed mutagenesis for investigating the activation mechanisms of ORs. The analysis of SlitOR25 and SlitOR31 binding cavities highlighted key residues, aligned with those found in the experimental MhraOR5^3^, ApisOR5^5^, AaegOR10^6^, and DmelOrco^65^ structures (Supplementary Table 4), that could be critical for the ligand binding and the activation mechanism. Moreover, the analysis of ligand-receptor interactions shows that in most of the cases SlitOR25 forms fewer hydrogen bonds with agonists than SlitOR31. This difference correlated with the higher polarity scores for the binding site of SlitOR31 compared to SlitOR25. Additionally, the cavity volume of SlitOR31 slightly exceeded that of SlitOR25 (roughly 530 vs 300 A^3^). It is consistent with the average number of heavy atoms (12 vs 8.4±1.4) and average size of SlitOR31 and SlitOR25 ligands (165±4 vs 123±13 A^3^). Our hypothesis is that the recognition spectrum of the receptor is partly driven by the size and polarity of the binding site. To support our findings (especially because of the limited number of SlitOR31 ligands), we analyzed the binding pockets of the 17 SlitORs from de Fouchier *et al*.^20^ Since all of these receptors were tested with the same 51 odorants, this provides clues about their tuning, enabling us to elaborate on structure-function relationship hypotheses. As shown in Supplementary Fig. 5 and Supplementary Table 5, the size and hydrophobicity of the binding pocket are correlated to the number of active molecules per receptor. Even if the correlation is somehow modest (Pearson’s r∼0.4-0.6), the larger the volume (or the solvent accessible surface area - SASA - of the pocket) and the lower the hydrophobicity, the narrower is the recognition spectrum of the receptor. This observation aligns with findings for SlitOR25 and SlitOR31, suggesting that such a tendency might be extrapolated to other insect ORs. We also attempted to build a simple model to predict the recognition spectrum of SlitORs. We set up a multiple linear regression model based on different combinations of the binding pocket descriptors to predict the experimental number of active molecules. As shown in Supplementary Fig. 6 and Supplementary Table 6, a very simple model with only two variables, SASA and hydrophobicity, performed well with a correlation coefficient of 0.75 between the experimental and the predicted values and a RMSE of 2.53. It already captured a large part of the structure-function relationship and was not much less powerful than the model with a higher number of variables (with 6 pocket descriptors, r=0.79 and RMSE=2.35). However, these conclusions must be put into perspective in view of the small size of the dataset. Moreover, a recent study on an OR-related sugar receptor highlights that chemical specificity is not exclusively determined by the selectivity of the ligand-binding pocket, but rather derives from multiple factors, primarily receptor-ligand interactions and allosteric coupling^73^. Anyhow, such a binding pocket analysis could guide the prediction of orphan receptors’ broadness.

The two new SlitOR25 ligands we identified in this study were attractants for *S. littoralis* larvae, as were the previously identified ligands^16,61^. In addition to reaffirming the effectiveness of reverse chemical ecology approaches, this discovery supports the hypothesis that activation of SlitOR25 induces larval attraction in *S. littoralis*. To our knowledge, the two new ligands have never been investigated in *S. littoralis* chemical ecology studies, and their ecological relevance can be questioned. Benzyl formate (16) induced the highest firing rate in OSNs expressing SlitOR25 and demonstrated a strong attraction rate for the larvae. Interestingly, this compound has been reported in volatile emissions of plant species within the genus *Solanum*: *S. stuckertii* and *S. incisum*^74^. As this plant genus includes the major host plants of *S. littorali*s, such as *S. lycopersicum*, *S. melongena* and *S. tuberosum*^75^, this cue may indicate the presence of suitable food sources for the larvae if emitted as well by these latter species. 3-(3,4- dihydroxyphenyl)propanoic acid (10) was the second compound we identified as activating SlitOR25 and inducing attraction of *S. littoralis* larvae. This molecule is present in various plant species, spanning herbs, shrubs and trees from distantly related families^76^. Interestingly, none of these plant species are known hosts of *S. littoralis*^75^, although some are present in the insect’s natural environment. As plant volatiles are constituted by a diversity of molecules, it is possible that other volatiles than 3-(3,4- dihydroxyphenyl)propanoic, detected by other SlitORs than OR25, disrupt the larvae attraction to this compound, leading to avoidance. Alternatively, unpalatable molecules may refrain larvae from feeding on these plants. The behavioral effect of the newly discovered ligand for SlitOR31, 2-methoxy-4- propylphenol (23), has not been tested in the current study since eugenol (the only previously identified ligand) is known to be inactive on *S. littoralis* larvae behavior^60^. However, eugenol has been shown to have antifeedant activity in larvae, resulting in their death if no other food source is present^77^. The close structural similarity between eugenol and 2-methoxy-4-propylphenol, only differing by the presence or absence of a double bond at the end of the aliphatic chain, suggests that both compounds may have similar effects on *S. littoralis*. This hypothesis is supported by a study demonstrating that applying 100 µg of 2-methoxy-4-propylphenol to a piece of maize significantly reduces feeding activities of larvae from the related species *S. frugiperda*^78^. A literature survey identified this compound in plant defense emissions of the oak *Quercus agrifolia*^79^. Whether it is also produced in non-host plants of *S. littoralis* remains to be investigated. Avoiding these cues could indeed enhance larval survival chances.

Altogether, our study paves the way for future research aimed at expanding the insect OR chemical space, which defines a given species’ olfactory capacities and shapes its chemical ecology. A comprehensive analysis of complete OR chemical spaces in both closely and distantly related species will ultimately unravel the intricate mechanisms of OR functionality and specialization and help us understand how OR evolution contributes to species adaptation to specific ecological niches. The Structure-Based Virtual Screening approach described here also opens new avenues for pest control via olfactory disruption, as it will undoubtedly accelerate the discovery of new behaviorally active volatiles.

## Acknowledgements

The authors thank Thomas Chertemps for insightful comments and fruitful discussions. This work was funded by the interdisciplinary MITI CNRS program and the French National Research Agency (ANR-20-CE20-003). This work was supported by the French government through the France 2030 investment plan managed by the National Research Agency (ANR), as part of the Initiative of Excellence Université Côte d’Azur under reference number ANR-15-IDEX-01. The authors are grateful to the Université Côte d’Azur’s Center for High-Performance Computing (OPAL infrastructure) for providing resources and support. The funders had no role in study design, data collection and analysis, decision to publish or preparation of the manuscript.

## Contributions

E.J.J and S.F. designed research; A.C, M.L., L.B. and R.M. performed research; A.C. analyzed data; A.C., E.J.J. and S.F. wrote the paper. All authors discussed the results, reviewed and approved the manuscript.

