## Supplementary Information for "Accelerating Ligand Discovery for Insect Odorant Receptors"

### Contents

**Supplementary Table 1:** List of compounds tested during electrophysiological assays and known ligands of SlitOR25 and SlitOR31.

**Supplementary Figure 1:** The binding sites of the 17 SlitORs models examined in this study, generated by either AlphaFold 2 or 3, exhibit a perfect match with the one identified in the experimental structure of MhraOR5.

**Supplementary Table 2:** LE stands out as the top-performing scoring function for analyzing the odorant docking poses on non-pheromonal odorant receptors of *Spodoptera littoralis*, including SlitOR25 and SlitOR31.

**Supplementary Figure 2:** The predicted chemical space of SlitOR25 and SlitOR31 is divided into 68 and 70 molecular clusters, respectively.

**Supplementary Table 3:** Performance of the *in silico* prediction for SlitOR25 and SlitOR31.

**Supplementary Table 4:** Summary of the residues in the binding pockets of SlitOR25 and SlitOR31 that were aligned with amino acids in the binding pockets of DmelOrco[2], MhraOR5[3], ApisOR5[5] and AaegOR10[6], where mutations have been shown to significantly impact ligand-receptor interactions.

**Supplementary Figure 3:** Effect of molecular weight and chemical family classification on the performance of Vinardo and LE scoring functions.

**Supplementary Figure 4:** Effect of the p-value threshold on the performance of the *in silico* prediction.

**Supplementary Figure 5:** Correlation between pocket descriptors and OR tuning.

**Supplementary Table 5:** Correlation matrix.

**Supplementary Figure 6:** Linear regression models for the prediction of OR broadness based on the description of the binding pocket.

**Supplementary Table 6:** Performance of the linear regression models.

**Supplementary Table 1:** List of compounds tested during electrophysiological assays and known ligands of SlitOR25 and SlitOR31. New ligands of SlitOR25 and SlitOR31 discovered with this present study are highlighted in grey.

The Supp. Table 1 is provided in the *Supplementary_Table1.xls* file

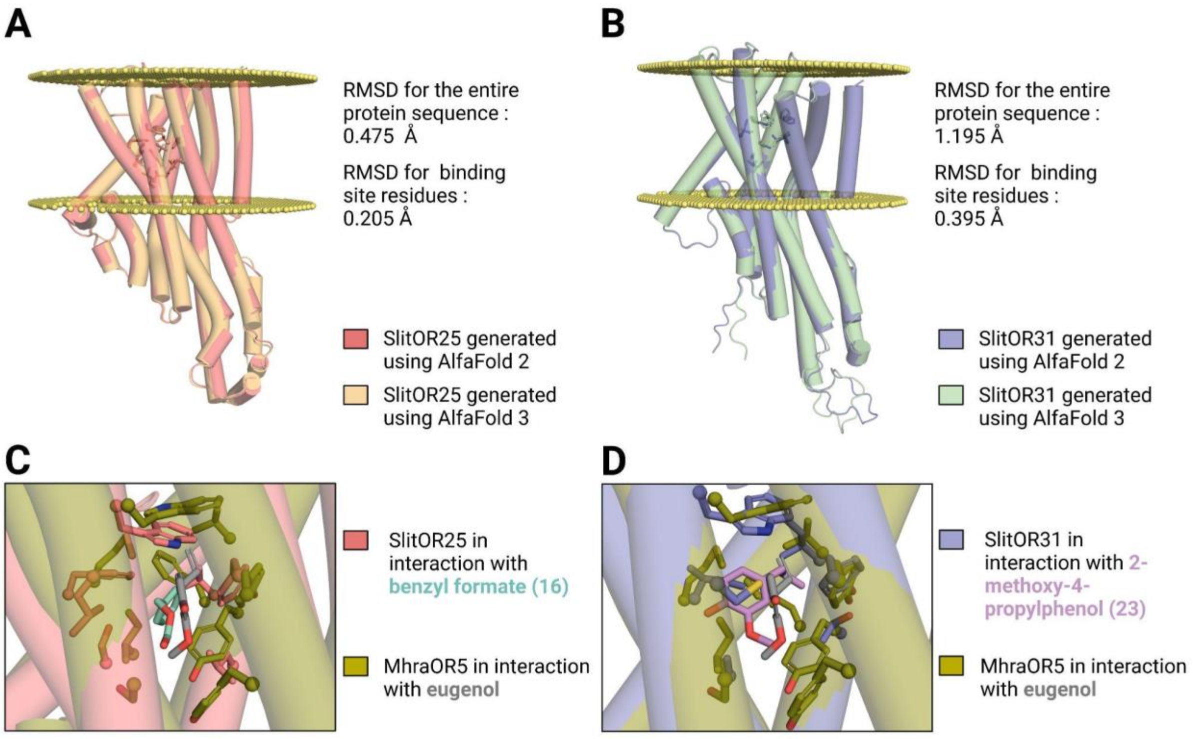


**Supplementary Figure 1: The binding sites of the 17 SlitORs models examined in this study, generated by either AlphaFold 2 or 3, exhibit a perfect match with the one identified in the experimental structure of MhraOR5.** Comparison of the structures generated by AlphaFold2 and AlphaFold3 for (**A**) SlitOR25 and (**B**) SlitOR31. (**C**) Superposition of the binding site from AlphaFold2 models of SlitOR25 and (**D**) SlitOR31 with the experimental structure of the MhraOR5, which binds eugenol. SlitOR25 is shown interacting with benzyl formate (16), while SlitOR31 interacts with 2-methoxy-4-propylphenol (23). The receptors orientation in the membrane was determined using OPM server[64]. This figure was generated using the molecular visualization software PyMol[48].

| | Max_EF_Vinardo | Threshold_Vinardo (%) | Max_EF_LEln | Threshold_LEln (%) | Max_EF_LESA | Threshold_LESA (%) | Max_EF_LE | Threshold_LE (%) |
|---|---|---|---|---|---|---|---|---|
| **Pheromone Receptors** | | | | | | | | |
| SlitOR6 | 5.1 | 20 | 3.92 | 25 | 1.31 | 76 | 1.11 | 90 |
| SlitOR13 | 12.75 | 4 | 12.75 | 8 | 2.68 | 37 | 1.46 | 69 |
| **Median** | **8.93** | **12** | **8.34** | **17** | **2.00** | **57** | **1.29** | **80** |
| **Non-pheromonal Receptors** | | | | | | | | |
| SlitOR3 | 2.32 | 4 | 2.32 | 8 | 1.45 | 63 | 1.55 | 53 |
| SlitOR4 | 1.16 | 86 | 3.4 | 6 | 5.10 | 4 | 10.20 | 2 |
| SlitOR7 | 1.06 | 94 | 1.04 | 96 | 1.38 | 73 | 2.27 | 29 |
| SlitOR14 | 1.11 | 90 | 1.34 | 75 | 4.25 | 12 | 12.75 | 2 |
| SlitOR17 | 1.89 | 53 | 1.82 | 55 | 2.55 | 16 | 6.80 | 6 |
| SlitOR19 | 1.09 | 92 | 1.21 | 82 | 2.12 | 31 | 6.38 | 8 |
| SlitOR24 | 1.16 | 86 | 1.76 | 57 | 5.67 | 2 | 5.67 | 2 |
| SlitOR25 | 1.15 | 80 | 1.57 | 59 | 2.24 | 14 | 3.92 | 2 |
| SlitOR27 | 1.00 | 100 | 1.00 | 100 | 1.14 | 63 | 2.43 | 6 |
| SlitOR28 | 1.06 | 94 | 1.11 | 90 | 1.28 | 78 | 2.43 | 14 |
| SlitOR29 | 1.24 | 80 | 1.11 | 90 | 2.43 | 59 | 2.02 | 35 |
| SlitOR31 | 1.34 | 75 | 2.04 | 49 | 5.10 | 10 | 8.50 | 6 |
| SlitOR32 | 1.09 | 92 | 1.19 | 84 | 1.42 | 47 | 5.67 | 2 |
| SlitOR35 | 1.75 | 24 | 2.14 | 27 | 3.00 | 2 | 3.00 | 2 |
| SlitOR36 | 2.83 | 18 | 4.25 | 23 | 4.63 | 22 | 8.50 | 2 |
| **Median** | **1.16** | **86** | **1.57** | **59** | **2.43** | **22** | **5.67** | **6** |

**Supplementary Table 2: LE stands out as the top-performing scoring function for analyzing the odorant docking poses on non-pheromonal odorant receptors of *Spodoptera littoralis*, including SlitOR25 and SlitOR31.** The comparison table summarizes the maximum enrichment factor values (Max_EF) obtained for the indicated threshold using Vinardo, LESA, LEln, and LE scoring functions based on optimal poses of 51 molecules across 17 receptors from de Fouchier et al., 2017[20]. The LESA, LEln, and LE functions are weighted versions of the Vinardo function as described in methodology. The gradient color scheme highlights AUC values, with green indicating higher performance and red lower ones.

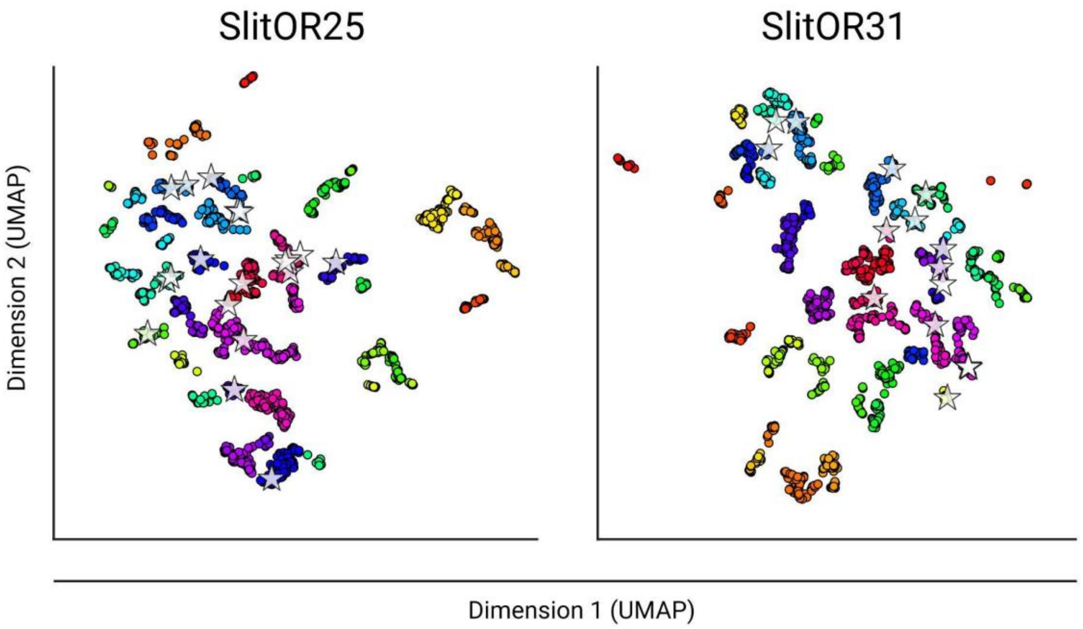


**Supplementary Figure 2: The predicted chemical space of SlitOR25 and SlitOR31 is divided into 68 and 70 molecular clusters, respectively.** UMAP representation of the chemical space based on Morgan2 fingerprints of potential agonists. The chemical space is divided into distinct clusters, each representing a group of potential agonists, with each cluster uniquely colored. Tested potential agonists are marked by white stars.

**Supplementary Table 3: Performance of the *in silico* prediction for** Slit**OR25 and** Slit**OR31.** P: Positives, N: Negatives, TP: True Positives, TN: True Negatives, FP: False Positives, FN: False Negatives.

| | Equations | SlitOR25 | SlitOR31 |
|---|---|---|---|
| **Sensitivity** | TPR = TP / (TP + FN) | 1.00 | 1.00 |
| **Specificity** | SPC = TN / (FP + TN) | 0.32 | 0.33 |
| **Precision** | PPV = TP / (TP + FP) | 0.11 | 0.06 |
| **Negative Predictive Value** | NPV = TN / (TN + FN) | 1.00 | 1.00 |
| **False Positive Rate** | FPR = FP / (FP + TN) | 0.68 | 0.67 |
| **False Discovery Rate** | FDR = FP / (FP + TP) | 0.89 | 0.94 |
| **False Negative Rate** | FNR = FN / (FN + TP) | 0.00 | 0.00 |
| **Accuracy** | ACC = (TP + TN) / (P + N) | 0.37 | 0.36 |
| **F1 Score** | F1 = 2TP / (2TP + FP + FN) | 0.19 | 0.11 |
| **Matthews Correlation Coefficient** | MCC = TP*TN - FP*FN / sqrt((TP+FP)*(TP+FN)*(TN+FP)*(TN+FN)) | 0.18 | 0.14 |

**Supplementary Table 4: Summary of the residues in the binding pockets of** Slit**OR25 and** Slit**OR31 that were aligned with amino acids in the binding pockets of DmelOrco[65], MhraOR5[3], ApisOR5[5] and AaegOR10[6], where mutations have been shown to significantly impact ligand-receptor interactions.** Colors indicate the impact of mutations on ligand recognition: grey highlights mutations that abolished the ligand recognition, blue denotes mutations that decreased receptor sensitivity, and orange indicates mutations that increased receptor sensitivity.

| SlitOR25 | SlitOR31 | MhraOR5 | DmelOrco | ApisOR5 | AaegOR10 |
|---|---|---|---|---|---|
| | F78 | | | | |
| V88 | I79 | V88D | | I63A | L67A |
| L91 | T82 | Y91A | F83A | | |
| Q92 | | F92A | F84A | | |
| | I141 | S151A | S146I | | |
| T153 | C144 | G154A | | | |
| L154 | G145 | | | | S133A |
| W157 | W148 | W158A | | F115A | F136A |
| F197 | I189 | M209A<br>M209L | V206W | V164A | |
| I200 | | | | | |
| S201 | A193 | I213A | | H168A | |
| V204 | | | | | |
| M205 | | | | | |
| F322 | Y313 | Y380A | | | |
| Y325 | | Y383A | | | |

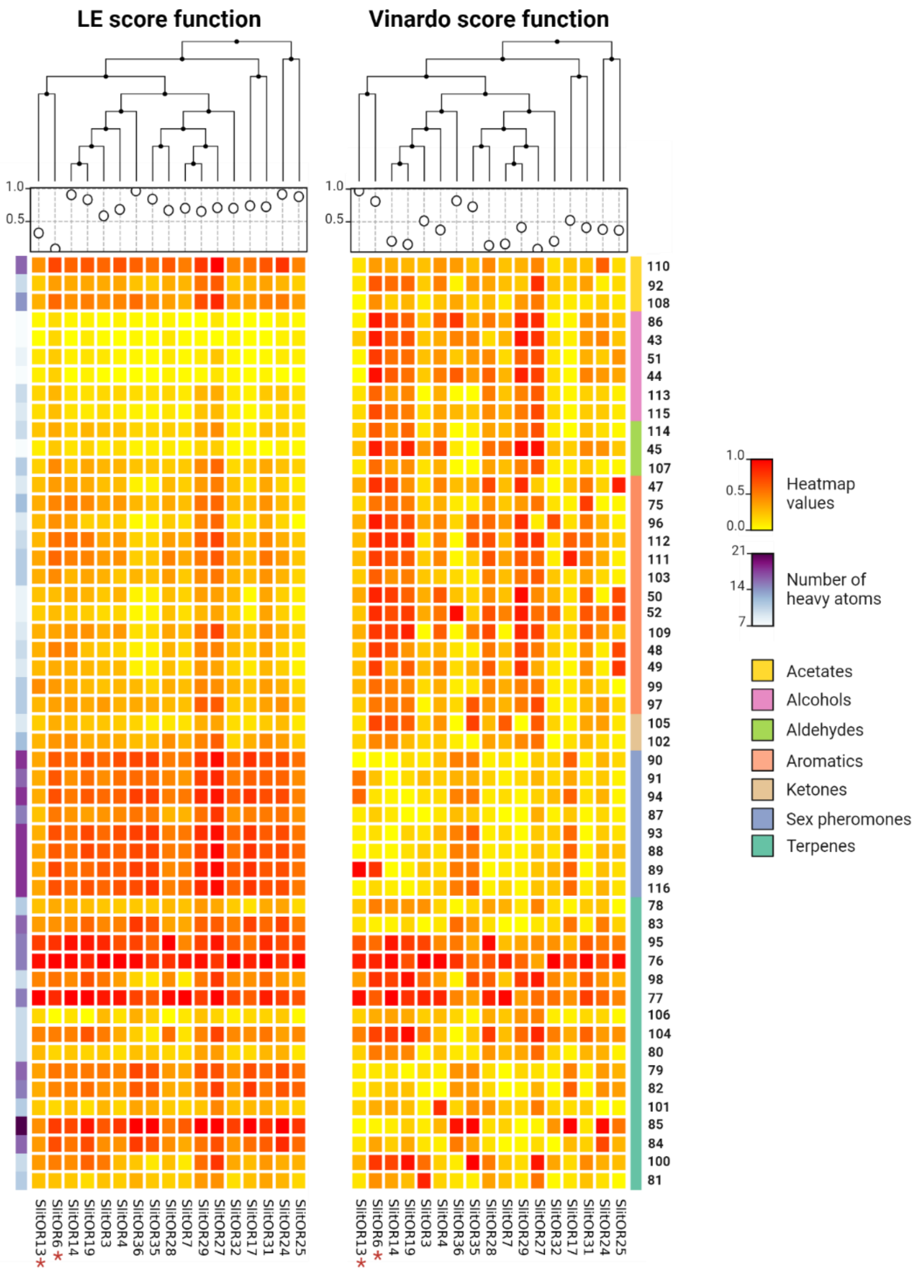


**Supplementary Figure 3: Effect of molecular weight and chemical family classification on the performance of Vinardo and LE scoring functions.** The two heatmaps depict the deviation between normalized experimental SSR values[20] and normalized docking values: the redder the color, the larger the deviation, indicating a greater difference between docking predictions and experimental values. Conversely, as the color tends towards yellow, docking values better align with experimental ones. The heatmap on the left was generated using the LE scoring function, while the one on the right was constructed using the unweighted Vinardo function. On the left side of the heatmaps, a blue gradient represents the number of heavy atoms for each molecule in rows: darker shades indicate higher atom counts. Above the heatmaps, a dot plot displays AUC values for all receptors in columns, and a dendrogram illustrates the clustering of these receptors. On the right of the heatmaps, the color code indicate the chemical families associated with each molecule: acetates in yellow, alcohols in pink, aldehydes in light green, aromatics in red, ketones in brown, sexual pheromones in blue, and terpenes in dark green. Molecules are labeled as in Supplementary Table 1. A red star denotes pheromone receptors.

**A**

| | LE ( p-value < 0.001) | LE (0.001< p-value < 0.01) | LE (0.01 < p-value < 0.05) |
|---|---|---|---|
| **Pheromone Receptors** | | | |
| SlitOR6 | 0.10 | 0.10 | 0.10 |
| SlitOR13 | 0.34 | 0.33 | 0.33 |
| **Median** | **0.22** | **0.22** | **0.22** |
| **Non-pheromonal Receptors** | | | |
| SlitOR3 | 0.59 | 0.67 | 0.80 |
| SlitOR4 | 0.69 | 0.69 | 0.69 |
| SlitOR7 | 0.71 | 0.65 | 0.75 |
| SlitOR14 | 0.91 | 0.85 | 0.84 |
| SlitOR17 | 0.75 | 0.84 | 0.84 |
| SlitOR19 | 0.83 | 0.86 | 0.84 |
| SlitOR24 | 0.92 | 0.92 | 0.85 |
| SlitOR25 | 0.87 | 0.87 | 0.87 |
| SlitOR27 | 0.72 | 0.77 | 0.81 |
| SlitOR28 | 0.68 | 0.79 | 0.77 |
| SlitOR29 | 0.66 | 0.67 | 0.57 |
| SlitOR31 | 0.72 | 0.72 | 0.82 |
| SlitOR32 | 0.71 | 0.74 | 0.74 |
| SlitOR35 | 0.85 | 0.90 | 0.89 |
| SlitOR36 | 0.97 | 0.98 | 0.93 |
| **Median** | **0.72** | **0.79** | **0.82** |

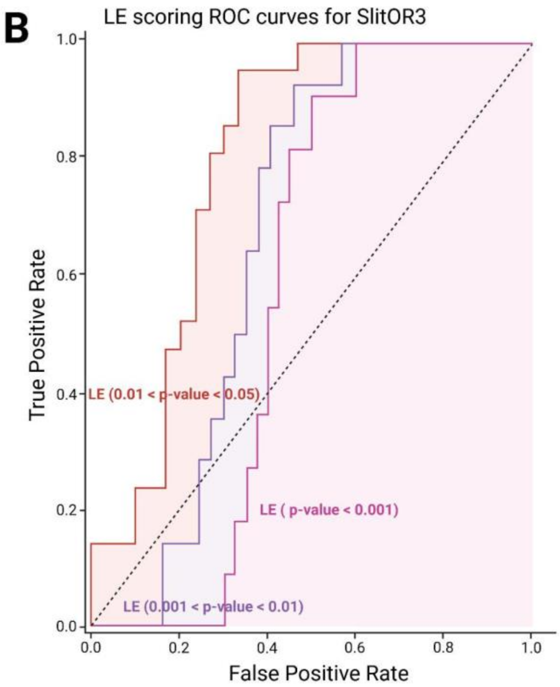


**Supplementary Figure 4 : Effect of the p-value threshold on the performance of the *in silico* prediction.** (**A**) The table summarizes AUC values from ROC curves using LE scoring function based on optimal poses of 51 molecules across 17 receptors from de Fouchier *et al.*, 2017[20]. Three different thresholds of p-values were considered to determine the activity of the molecule on the SlitORs from experimental data: p-value < 0.001; 0.001 < p-value < 0.01; and 0.01 < p-value < 0.05. The gradient color scheme highlights AUC values, with green indicating higher performance while red are lower ones. (**B**) ROC curves for SlitOR3 show performance with LE scoring function and a p-value < 0.001 (pink); 0.001 < p-value < 0.01 (purple); and 0.01 < p-value < 0.05 (orange).

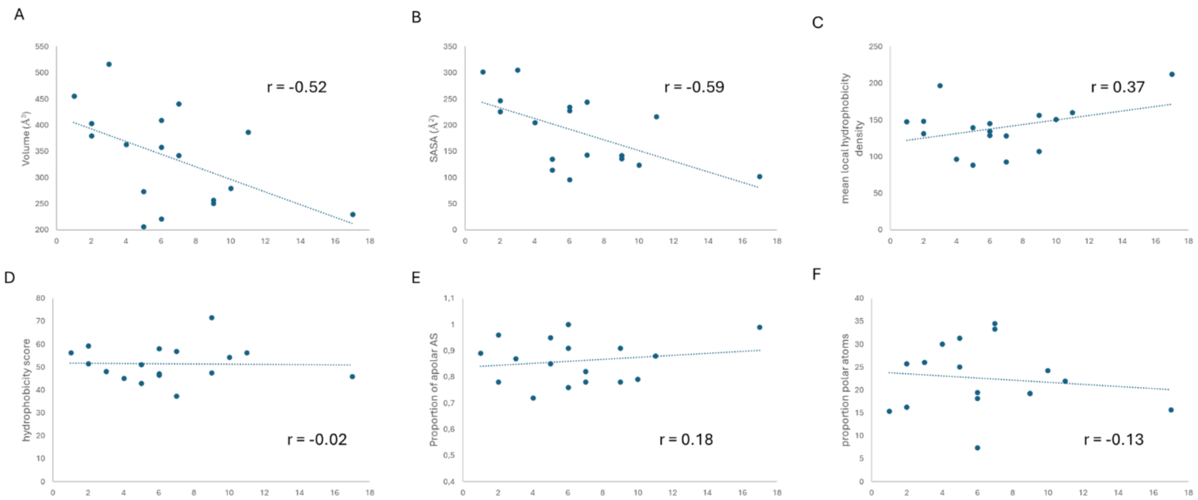


**Supplementary Figure 5: Correlation between pocket descriptors and OR tuning.** We have retained the descriptors of the binding cavity that are most relevant from a statistical standpoint (those with the best correlation to the number of ligands per OR) and for their ability to facilitate the deduction of structure-function relationships. The 6 descriptors retained for analysis are as follows: (**A**) Pocket volume, (**B**) Total surface area, (**C**) Mean local hydrophobic density, (**D**) Hydrophobicity score, (**E**) Proportion of apolar alpha spheres and (**F**) Proportion of polar atoms. In the analysis, we did not retain the Volume score, Charge score and Polarity Score as, based on the author comments, these descriptors are extremely approximative and should not be overestimated. The descriptors describing the surface area of the binding pocket (pock_pol_asa, pock_apol_asa, ...) were not significant and too dependent on the absolute value of the Total surface area of the pocket. We decided to calculate a relative value, i.e. the ratio between the apolar surface area and the total surface area. This ratio is significantly correlated to other descriptors (proportion of apolar alpha sphere, Mean local hydrophobic density) already used for the analysis and was finally not kept. The descriptors describing the alpha spheres were also not considered in the analysis as they do not provide a satisfactory explanation and are not easily interpretable. Finally, the amino acid composition of the cavity does not provide a satisfactory explanation, probably due to the limited size of the dataset (17 OR).

**Supplementary Table 5: Correlation matrix.** Pearson's correlation coefficient between the number of active ligands (nb_ligands) and pocket descriptors calculated with MDpocket. One has to note that SASA and Volume are highly correlated (r >0.9). To avoid bias in the analysis, we decided to reduce to only 2 variables the linear models (mean_Hyd_dens + SASA or Volume).

| | | nb_ligands | Volume | SASA | AS_apol_prop | mean_Hyd_dens | Hyd_scor | prop_polar_atm |
|---|---|---|---|---|---|---|---|---|
| **Volume** | *Pearson's r* | -0.516 | — | | | | | |
| | *p-value* | 0.034 | — | | | | | |
| **SASA** | *Pearson's r* | -0.591 | 0.941 | — | | | | |
| | *p-value* | 0.013 | < .001 | — | | | | |
| **AS_apol_prop** | *Pearson's r* | 0.180 | -0.345 | -0.280 | — | | | |
| | *p-value* | 0.490 | 0.176 | 0.276 | — | | | |
| **mean_Hyd_dens** | *Pearson's r* | 0.367 | 0.187 | 0.158 | 0.368 | — | | |
| | *p-value* | 0.147 | 0.473 | 0.545 | 0.146 | — | | |
| **Hyd_scor** | *Pearson's r* | -0.021 | -0.103 | 0.029 | -0.098 | -0.183 | — | |
| | *p-value* | 0.937 | 0.695 | 0.911 | 0.709 | 0.483 | — | |
| **prop_polar_atm** | *Pearson's r* | -0.125 | 0.265 | 0.080 | -0.462 | -0.371 | -0.290 | — |
| | *p-value* | 0.634 | 0.303 | 0.760 | 0.062 | 0.142 | 0.259 | — |

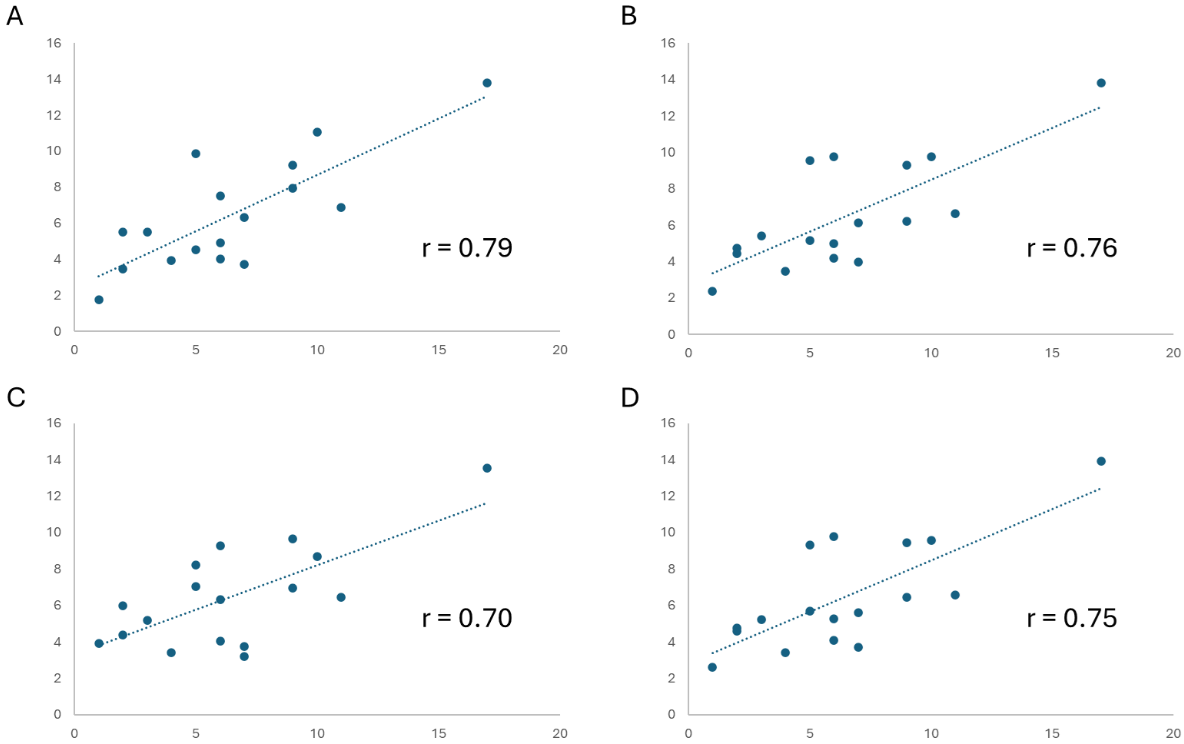


**Supplementary Figure 6: Linear regression models for the prediction of OR broadness based on the description of the binding pocket.** The scatter plot represents the predicted *vs* experimental predicted number of ligands per OR for each model. **(A)** The prediction is based on a multiple linear regression (MLR) model including the 6 normalized descriptors as described in the Supplementary Figure 5. (**B**) MLR includes only 3 descriptors, the most significant: Pocket volume, Total surface area and Mean local hydrophobic density. (**C**) MLR includes only 2 descriptors: Volume and Mean local hydrophobic density and (**D**) same as C with Total surface area and Mean local hydrophobic density.

The equation of the model in D) is *NL = -8.07 * SASA + 6.98 * Hyd_dens + 7.18* where *NL* means the predicted number of ligands per OR, *SASA* is the total surface area of the binding pocket and *Hyd_dens* is the mean local hydrophobic density. For this model, Pearson's r=0.75 and RMSE=2.53.

**Supplementary Table 6: Performance of the linear regression models.** As expected, the model with the higher number of variables yields the best performance. Since three descriptors are too poorly correlated (See Supp. Table 3) and the pocket SASA is highly correlated to the volume, the model with only 2 descriptors reaches similar performance.

| Models | r | $r^2$ | RMSE |
|---|---|---|---|
| 6 desc | 0.79 | 0.63 | 2.35 |
| 3 desc | 0.76 | 0.57 | 2.51 |
| 2 desc* | 0.70 | 0.49 | 2.74 |
| 2 desc** | 0.75 | 0.57 | 2.53 |

* The 2 descriptors used in the model are Volume and Mean hydrophobicity density
** The 2 descriptors used in the model are SASA and Mean hydrophobicity density
